# Comprehensive in silico Analysis of *IKBKAP* gene that could potentially cause Familial dysautonomia

**DOI:** 10.1101/436071

**Authors:** Mujahed I. Mustafa, Enas A. Osman, Abdelrahman H. Abdelmoneiom, Dania M. Hassn, Hadeel M. Yousif, Inshrah K. Mahgoub, Razan M. Badawi, Kutuf A. Albushra, Tebyan A Abdelhameed, Mohamed A. Hassan

**Affiliations:** Department of Biochemistry, University of Bahri, Sudan; Department of Biotechnology, Africa city of Technology, Sudan

**Author notes:** Crossponding author: Mujahed I. Mustafa.

**Keywords:** Familial dysautonomia (FD), neurodevelopmental genetic disorder, autonomic neuropathies, *SNPs*, *computational analysis*, diagnostic markers

## Abstract

**Background:** Familial dysautonomia (FD) is a rare neurodevelopmental genetic disorder within the larger classification of hereditary sensory and autonomic neuropathies. *We aimed to identify the pathogenic SNPs in IKBKAP* gene *by computational analysis software’s, and to determine the structure, function and regulation of their respective proteins.*

**Materials and Methods:** *We carried out in silico analysis of structural effect of each SNP using different bioinformatics tools to predict SNPs influence on protein structure and function.*

**Result:** *41 novel mutations out of 973 nsSNPs that are found be deleterious effect on the IKBKAP structure and function.*

**Conclusion:** *This is the first in silico analysis in IKBKAP gene to prioritize SNPs for further genetic studies.*

## Introduction

Familial dysautonomia (FD; also known as “Riley-Day syndrome”) is a rare neurodevelopmental genetic disorder within the larger classification of hereditary sensory and autonomic neuropathies,^(1, 2) (3) (4) (5)^ each caused by a different genetic error associated with an increased risk for sudden death.^(6, 7) (8)^ it is also called hereditary sensory and autonomic neuropathy type III (HSAN III) which affects 1/3600 live births in the Ashkenazi Jewish population.^(9)^

The *IKBKAP* gene give specific instructions for making a protein called elongator complex protein-1 (ELP1). This protein is found in a variety of cells, including brain cells. It is part of a six-protein complex called the elongator complex. The elongator complex plays a key role in transcription.^(10, 11)^ The disease is caused by mutation in the *IKBKAP* (*ELP1*) gene ^(8, 12)^ that affects the splicing of the elongator-1 protein (ELP-1) (also known as IKAP). But how it functions in disease-vulnerable neurons is still unknown. The mutation weakens the 5′ splice site of exon 20. Variable skipping of exon 20 leads to a tissue-specific reduction in the level of ELP1 protein. Which has been mapped to a 0.5-cM region on chromosome 9q31, has eluded identification.^(13, 14)^

The most common mutation, which is present on 99.5% of all FD chromosomes, is an intronic splice site mutation that results in tissue-specific skipping of exon 20. The second FD mutation, a missense change in exon 19 (R696P), is a missense mutation that has been identified in 4 unrelated patients heterozygous for the major splice mutation. Interestingly, despite the fact that FD is a recessive disease, normal mRNA and protein are expressed in patient cells, the diagnosis of FD has been limited to individuals of Ashkenazi Jewish ^(5)^ descent and identification of the gene has led to widespread diagnostic and carrier testing in this population. The first non-Jewish IKBKAP mutation was reported, a proline to leucine missense mutation in exon 26, (P914L). This mutation is of particular significance because it was identified in a patient who lacks one of the cardinal diagnostic criteria for the disease-pure Ashkenazi Jewish ancestry.^(15, 16)^

The disorder disturbs cells in the autonomic nervous system, which controls involuntary actions such as digestion, breathing (patients develop neurogenic dysphagia with frequent aspiration, chronic lung disease, and chemoreflex failure leading to severe sleep disordered breathing.^(17)^ ,production of tears, and the regulation of blood pressure and body temperature. It also affects the sensory nervous system, which controls activities related to the senses. ^(4) (8) (18) (19)^ Progressive ataxic gait is also a common symptom in FD patients. At least 50% of adults with FD require assistance with walking. ^(20)^

The aim of this study was to identify the pathogenic SNPs in *IKBKAP* using in silico prediction softwares, and to determine the structure, function and regulation of their respective proteins. This is the first in silico analysis in *IKBKAP* gene to prioritize SNPs for further genetic mapping studies. The usage of in silico approach has strong impact on the identification of candidate SNPs since they are easy and less costly, and can facilitate future genetic studies. ^(21)^

## 2. Method

### 2.1 Data mining

The data on human *IKBKAP* gene was collected from National Center for Biological Information (NCBI) web site ^(22)^ The SNP information (protein accession number and SNP ID) of the *MEFV* gene was retrieved from the NCBI dbSNP (http://www.ncbi.nlm.nih.gov/snp/) and the protein sequence was collected from Swiss Prot databases (http://expasy.org/). ^(23)^

### 2.2 SIFT

SIFT is a sequence homology-based tool ^(24)^ that sorts intolerant from tolerant amino acid substitutions and predicts whether an amino acid substitution in a protein will have a phenotypic Effect. Considers the position at which the change occurred and the type of amino acid change. Given a protein sequence, SIFT chooses related proteins and obtains an alignment of these proteins with the query. Based on the amino acids appearing at each position in the alignment, SIFT calculates the probability that an amino acid at a position is tolerated conditional on the most frequent amino acid being tolerated. If this normalized value is less than a cutoff, the substitution is predicted to be deleterious. SIFT scores <0.05 are predicted by the algorithm to be intolerant or deleterious amino acid substitutions, whereas scores >0.05 are considered tolerant. It is available at (http://sift.bii.a-star.edu.sg/).

### 2.3 Polyphen-2

It is a software tool ^(25)^ to predict possible impact of an amino acid substitution on both structure and function of a human protein by analysis of multiple sequence alignment and protein 3D structure, in addition it calculates position-specific independent count scores (PSIC) for each of two variants, and then calculates the PSIC scores difference between two variants. The higher a PSIC score difference, the higher the functional impact a particular amino acid substitution is likely to have. Prediction outcomes could be classified as probably damaging, possibly damaging or benign according to the value of PSIC as it ranges from (0_1); values closer to zero considered benign while values closer to 1 considered probably damaging and also it can be indicated by a vertical black marker inside a color gradient bar, where green is benign and red is damaging. nsSNPs that predicted to be intolerant by Sift has been submitted to Polyphen as protein sequence in FASTA format that obtained from UniproktB /Expasy after submitting the relevant ensemble protein (ESNP) there, and then we entered position of mutation, native amino acid and the new substituent for both structural and functional predictions. PolyPhen version 2.2.2 is available at http://genetics.bwh.harvard.edu/pph2/index.shtml

### 2.4 Provean

Provean is a software tool ^(26)^ which predicts whether an amino acid substitution or indel has an impact on the biological function of a protein. It is useful for filtering sequence variants to identify nonsynonymous or indel variants that are predicted to be functionally important. It is available at (https://rostlab.org/services/snap2web/).

### 2.5 SNAP2

Functional effects of mutations are predicted with SNAP2 ^(27)^ SNAP2 is a trained classifier that is based on a machine learning device called “neural network”. It distinguishes between effect and neutral variants/non-synonymous SNPs by taking a variety of sequence and variant features into account. The most important input signal for the prediction is the evolutionary information taken from an automatically generated multiple sequence alignment. Also structural features such as predicted secondary structure and solvent accessibility are considered. If available also annotation (i.e. known functional residues, pattern, regions) of the sequence or close homologs are pulled in. In a cross-validation over 100,000 experimentally annotated variants, SNAP2 reached sustained two-state accuracy (effect/neutral) of 82% (at an AUC of 0.9). In our hands this constitutes an important and significant improvement over other methods. It is available at (https://rostlab.org/services/snap2web/).

### 2.6 PHD-SNP

An online Support Vector Machine (SVM) based classifier, is optimized to predict if a given single point protein mutation can be classified as disease-related or as a neutral polymorphism, it is available at: (http://snps.biofold.org/phd-snp/phdsnp.html)

### 2.7 SNP& Go

SNPs&GO is an algorithm developed in the Laboratory of Biocomputing at the University of Bologna directed by Prof. Rita Casadio. SNPs&GO is an accurate method that, starting from a protein sequence, can predict whether a variation is disease related or not by exploiting the corresponding protein functional annotation. SNPs&GO collects in unique framework information derived from protein sequence, evolutionary information, and function as encoded in the Gene Ontology terms, and outperforms other available predictive methods ^(28)^ It is available at (http://snps.biofold.org/snps-and-go/snps-and-go.html).

### 2.8 P-Mut

PMUT a web-based tool (29) for the annotation of pathological variants on proteins, allows the fast and accurate prediction (approximately 80% success rate in humans) of the pathological character of single point amino acidic mutations based on the use of neural networks. It is available at (http://mmb.irbbarcelona.org/PMut).

### 2.9 I-Mutant 3.0

I-Mutant 3.0 Is a neural network based tool ^(30)^ for the routine analysis of protein stability and alterations by taking into account the single-site mutations. The FASTA sequence of protein retrieved from UniProt is used as an input to predict the mutational effect on protein stability. It is available at (http://gpcr2.biocomp.unibo.it/cgi/predictors/I-Mutant3.0/I-Mutant3.0.cgi).

### 2.10 Modeling nsSNP locations on protein structure

Project hope (version 1.1.1) is a new online web-server to search protein 3D structures (if available) by collecting structural information from a series of sources, including calculations on the 3D coordinates of the protein, sequence annotations from the UniProt database, and predictions by DAS services. Protein sequences were submitted to project hope server in order to analyze the structural and conformational variations that have resulted from single amino acid substitution corresponding to single nucleotide substitution; It is available at (http://www.cmbi.ru.nl/hope)

### 2.11 GeneMANIA

We submitted genes and selected from a list of data sets that they wish to query. GeneMANIA approach to know protein function prediction integrate multiple genomics and proteomics data sources to make inferences about the function of unknown proteins. (31) . It is available at (http://www.genemania.org/).

### 2.12 RaptorX

is a web server predicting structure property of a protein sequence without using any templates. It outperforms other servers, especially for proteins without close homologs in PDB or with very sparse sequence profile. The server predicts tertiary structure. It is available at (http://raptorx.uchicago.edu/) (32)

## 3. Result

## Discussion

We found *41 novel mutations (table3) that has* effect on the stability and function of the *IKBKAP* gene using bioinformatics tools. The methods used are based on different aspects and parameters describing the pathogenicity and provide clues on the molecular level about the effect of mutations. It is not easy to predict the pathogenic effect of SNPs using single method. Therefore, we used multiple methods to compare and rely on the results predicted. In this study we used different in silico prediction algorithms: SIFT, PolyPhen-2, Provean, SNAP2, SNP&GO, PHD-SNP, P-MUT and I-Mutant 3.0 ,Figure (2).

This study identified the total number of nsSNP in Homo sapiens located in coding region of *IKBKAP* gene, that were investigated in dbSNP/NCBI database ^(22)^ out of 1458 there are 973 nsSNPs (missense mutations), which were submitted to SIFT server, PolyPhen-2 server, Provean sever and SNAP2 respectively, 291 (130 SNPs with Score 0) SNPs were predicted to be deleterious in SIFT server. In PolyPhen-2 server, our result showed that 570 were found to be damaging (159 possibly damaging and 411 probably damaging showed deleterious). In Provean server our result showed that 390 SNPs were predicted to be deleterious. While in SNAP2 server our result showed that 389 SNPs were predicted to be Effect. The differences in prediction capabilities refer to the fact that every prediction algorithm uses different sets of sequences and alignments. In table (2) we were submitted four positive results from SIFT, PolyPhen-2, Provean and SNAP2 to observe the disease causing one by SNP&GO, PHD-SNP and P-Mut servers. Figures (1 & 2).

**Figure 1:**
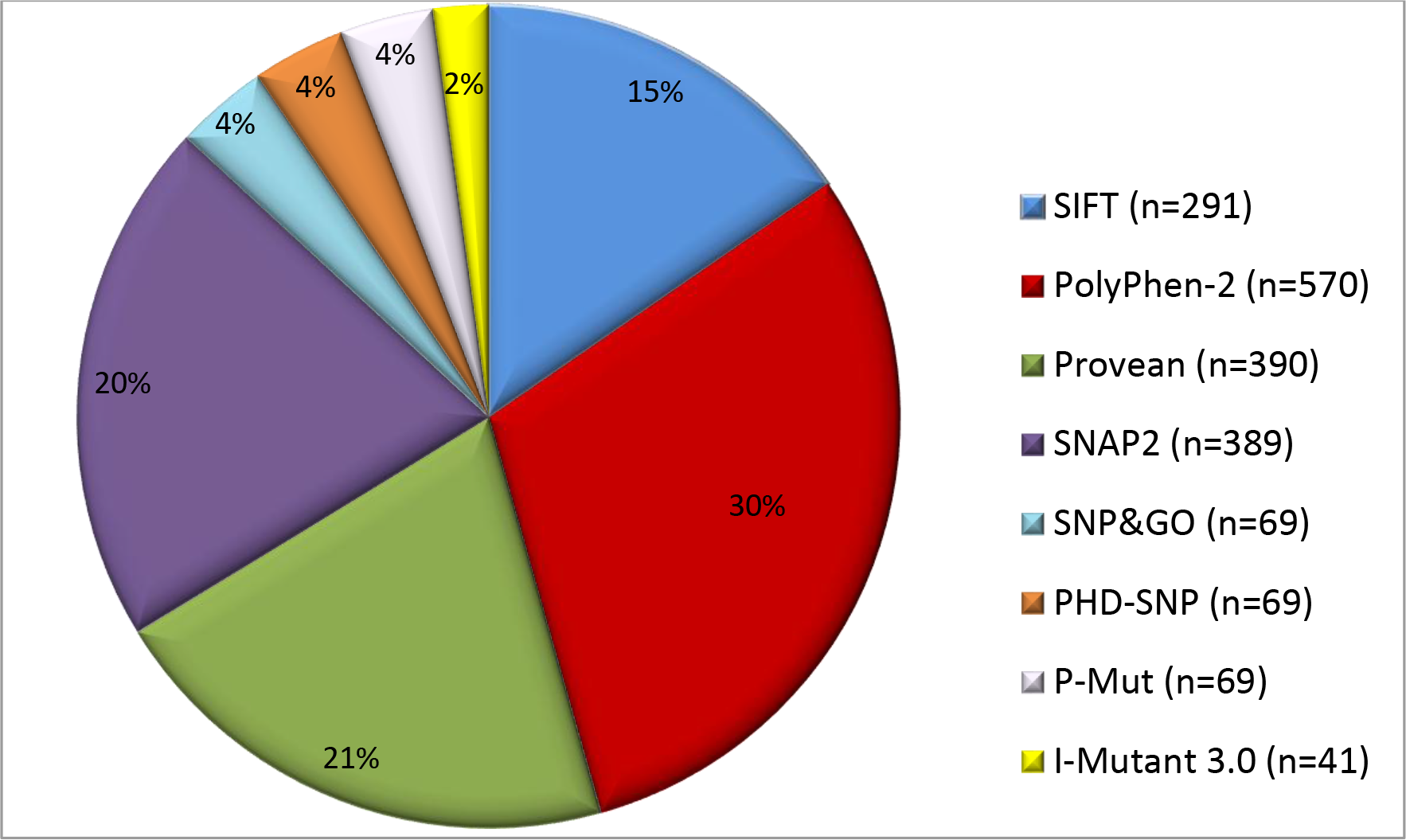
Pie chart showing percentage of damaging nsSNPs identified. The distribution of damaging nsSNPs by percentage (%) and number of SNPs (n) identified by eight in silico tools; SIFT, PolyPhen-2, Provean, SNAP2, SNP&GO, PHD-SNP, P-MUT and I-Mutant 3.0.

**Figure 2:**
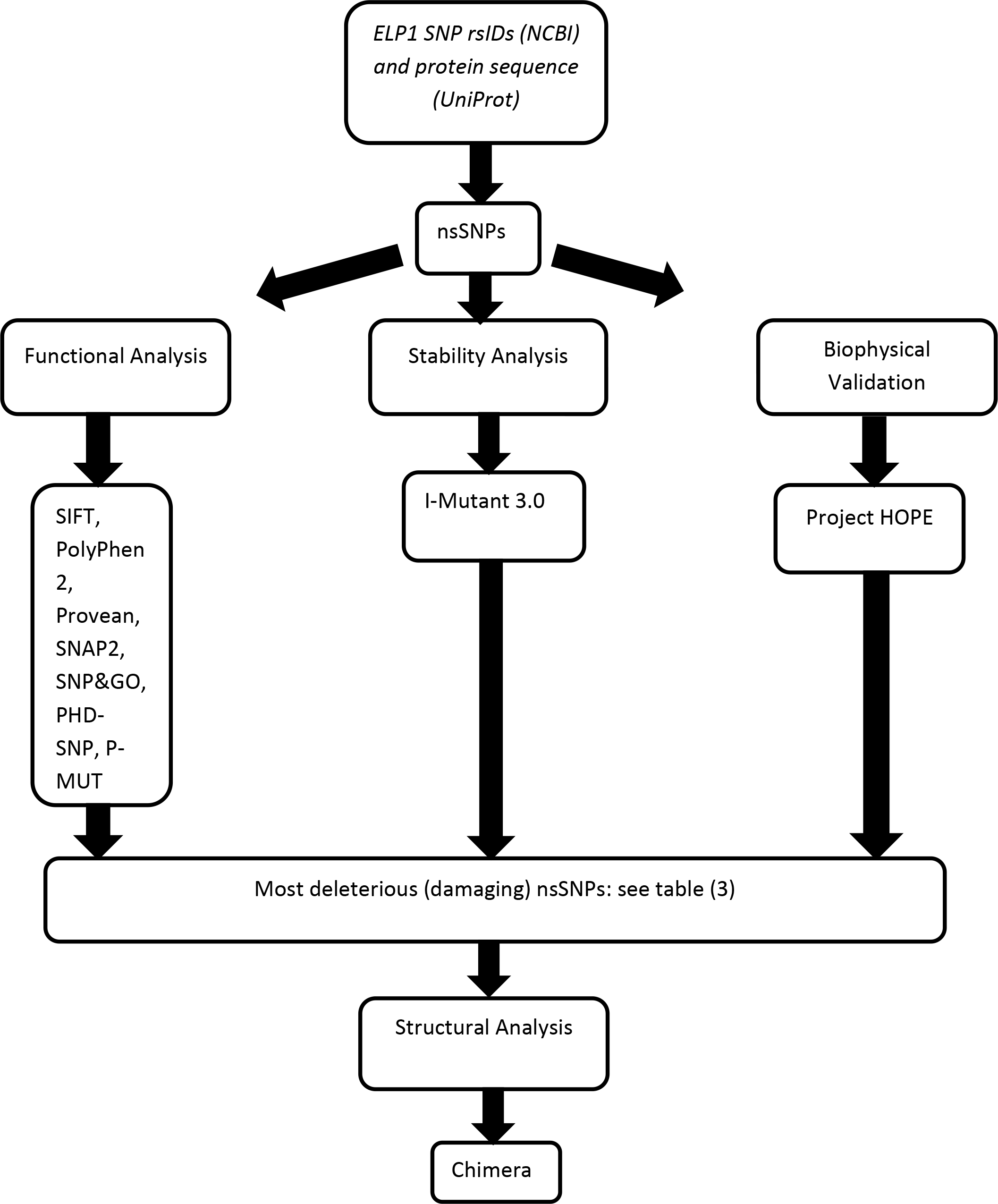
Diagrammatic representation of *IKBKAP* gene in silico work flow.

**Figure 3:**
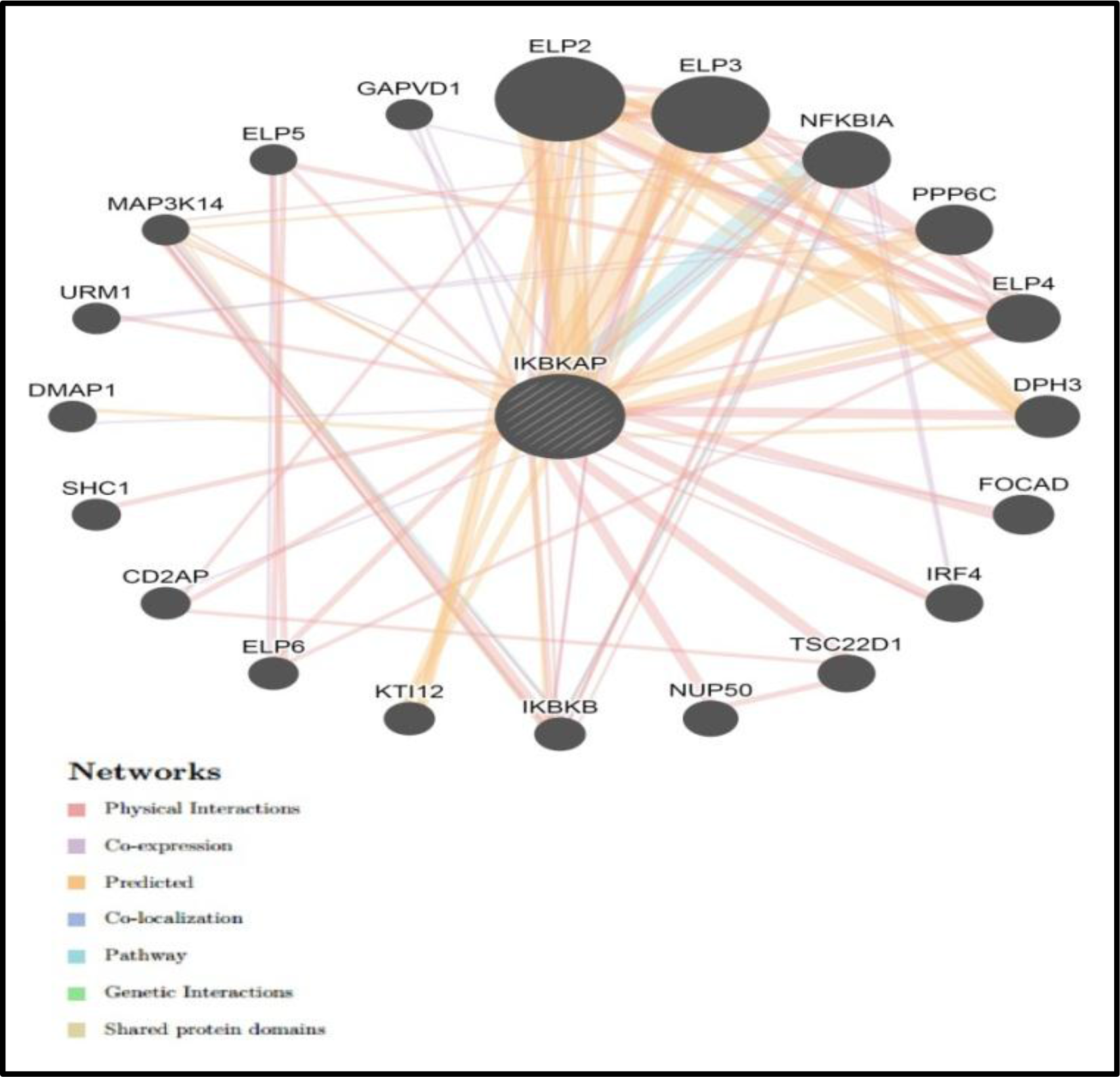
interaction between *IKBKAP* and its related genes.

**Table 1:**
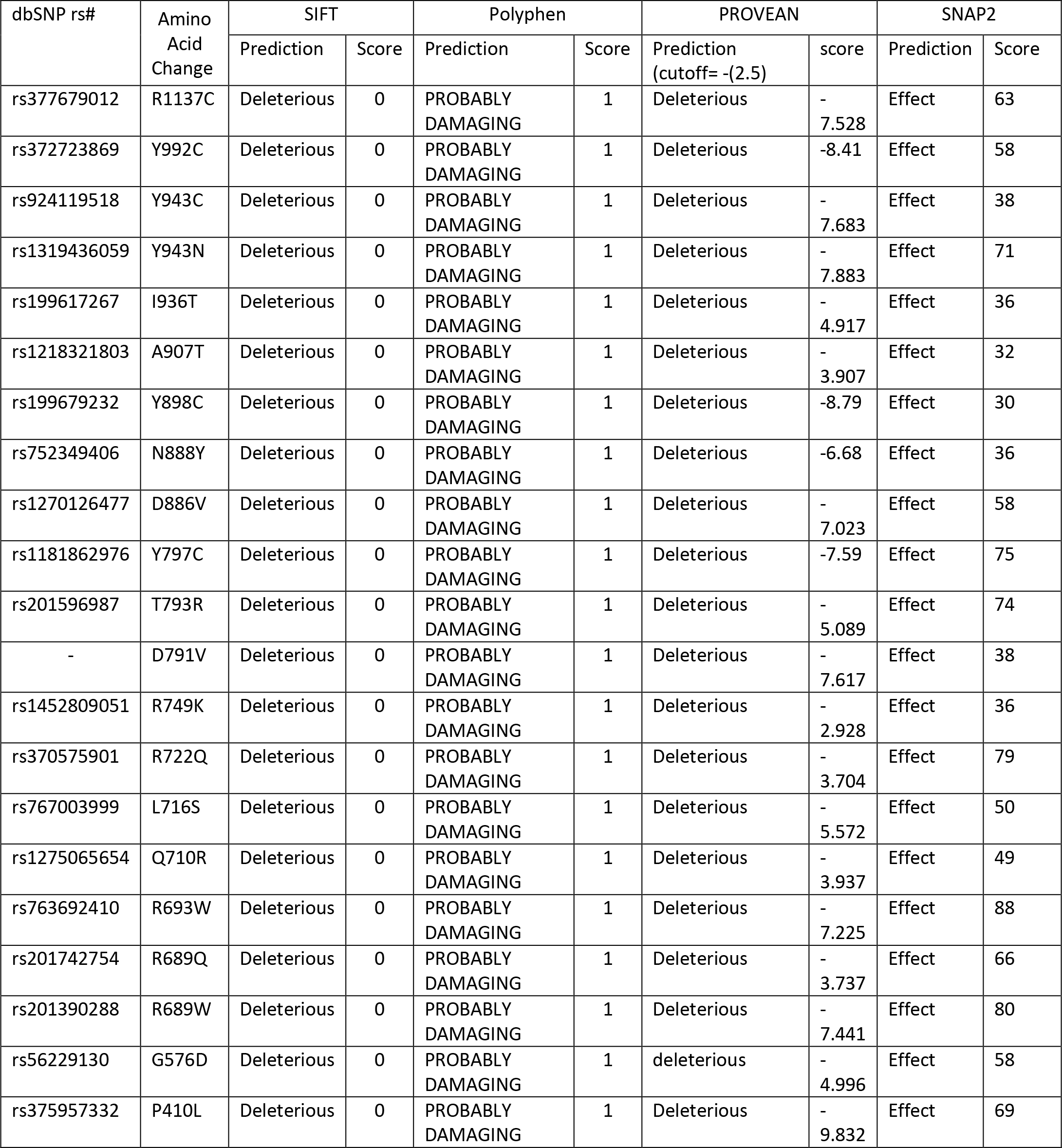
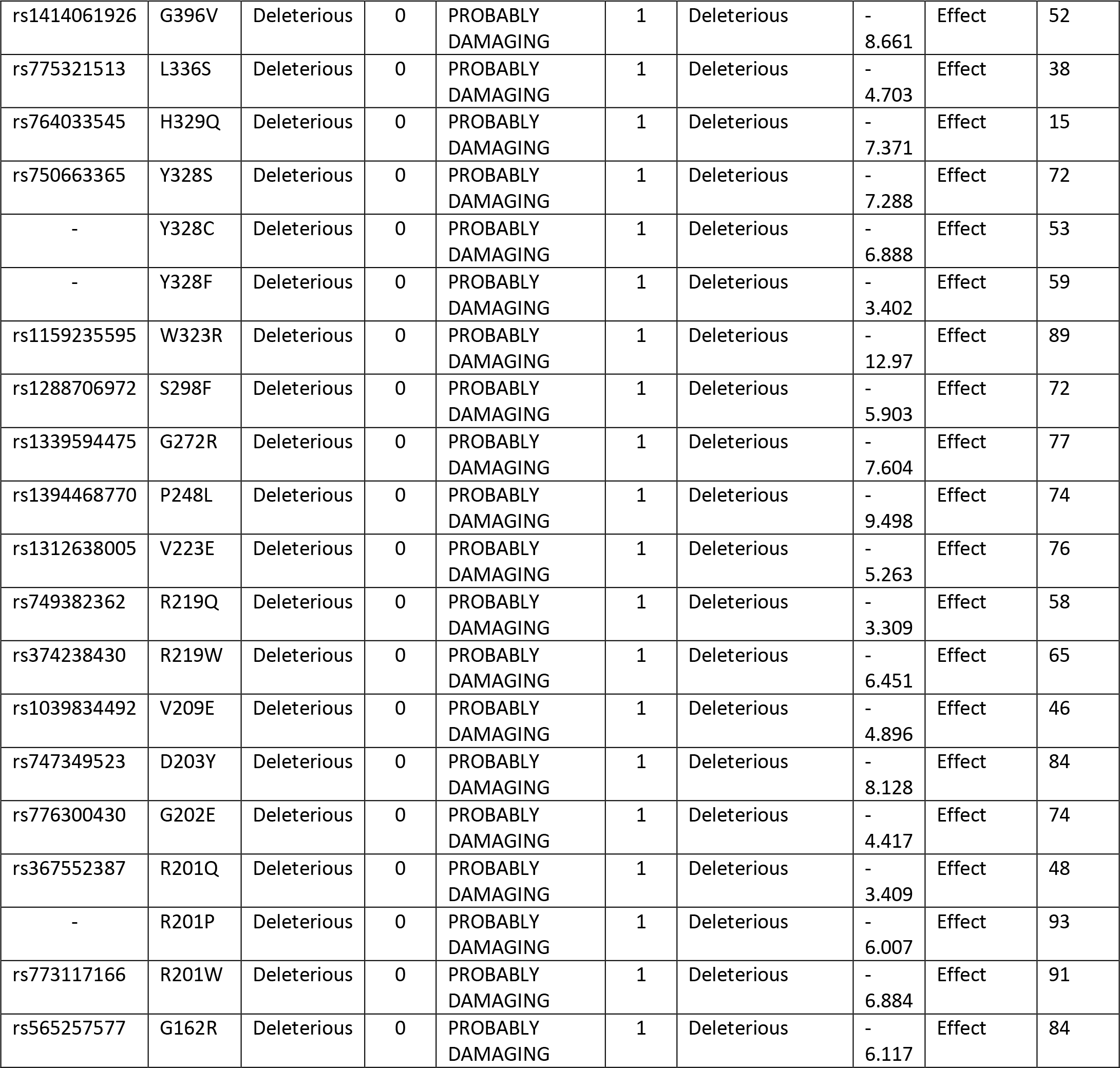
Damaging or Deleterious or effect nsSNPs associated variations predicted by various softwares:

**Table 2:** Disease effect nsSNPs associated variations predicted by various softwares:

| dbSNP rs# | Amino acid change | SNP&GO |  |  | PHD-SNP |  |  | P-Mut |  |
| --- | --- | --- | --- | --- | --- | --- | --- | --- | --- |
|  |  | Prediction | R I | Score | Prediction | R I | Score | Score | Prediction |
| rs377679012 | R1137C | Disease | 5 | 0.736 | Disease | 6 | 0.805 | 0.90 (93%) | Disease |
| rs372723869 | Y992C | Disease | 2 | 0.579 | Disease | 3 | 0.665 | 0.54 (80%) | Disease |
| rs924119518 | Y943C | Disease | 0 | 0.525 | Disease | 3 | 0.629 | 0.79 (89%) | Disease |
| rs1319436059 | Y943N | Disease | 2 | 0.583 | Disease | 4 | 0.687 | 0.70 (86%) | Disease |
| rs199617267 | I936T | Disease | 6 | 0.794 | Disease | 8 | 0.879 | 0.86 (91%) | Disease |
| rs1218321803 | A907T | Disease | 1 | 0.534 | Disease | 3 | 0.635 | 0.86 (91%) | Disease |
| rs199679232 | Y898C | Disease | 7 | 0.851 | Disease | 8 | 0.895 | 0.83 (90%) | Disease |
| rs752349406 | N888Y | Disease | 2 | 0.599 | Disease | 5 | 0.749 | 0.59 (82%) | Disease |
| rs1270126477 | D886V | Disease | 6 | 0.783 | Disease | 7 | 0.855 | 0.81 (89%) | Disease |
| rs1181862976 | Y797C | Disease | 4 | 0.686 | Disease | 5 | 0.742 | 0.86 (91%) | Disease |
| rs201596987 | T793R | Disease | 3 | 0.646 | Disease | 2 | 0.589 | 0.76 (88%) | Disease |
| - | D791V | Disease | 4 | 0.695 | Disease | 5 | 0.732 | 0.81 (89%) | Disease |
| rs1452809051 | R749K | Disease | 1 | 0.562 | Disease | 3 | 0.652 | 0.58 (82%) | Disease |
| rs370575901 | R722Q | Disease | 7 | 0.829 | Disease | 8 | 0.9 | 0.69 (85%) | Disease |
| rs767003999 | L716S | Disease | 5 | 0.757 | Disease | 7 | 0.848 | 0.81 (89%) | Disease |
| rs1275065654 | Q710R | Disease | 4 | 0.701 | Disease | 4 | 0.69 | 0.75 (87%) | Disease |
| rs763692410 | R693W | Disease | 3 | 0.63 | Disease | 8 | 0.899 | 0.86 (91%) | Disease |
| rs201742754 | R689Q | Disease | 4 | 0.707 | Disease | 7 | 0.872 | 0.70 (86%) | Disease |
| rs201390288 | R689W | Disease | 5 | 0.728 | Disease | 8 | 0.911 | 0.84<br>(90%) | Disease |
| rs56229130 | G576D | Disease | 6 | 0.821 | Disease | 7 | 0.87 | 0.60<br>(83%) | Disease |
| rs375957332 | P410L | Disease | 6 | 0.804 | Disease | 7 | 0.859 | 0.88<br>(92%) | Disease |
| rs141406192<br>6 | G396V | Disease | 6 | 0.819 | Disease | 9 | 0.932 | 0.86<br>(91%) | Disease |
| rs775321513 | L336S | Disease | 2 | 0.612 | Disease | 2 | 0.589 | 0.67<br>(85%) | Disease |
| rs764033545 | H329Q | Disease | 6 | 0.813 | Disease | 8 | 0.9 | 0.59<br>(82%) | Disease |
| rs750663365 | Y328S | Disease | 7 | 0.829 | Disease | 8 | 0.881 | 0.84<br>(90%) | Disease |
| - | Y328C | Disease | 7 | 0.834 | Disease | 8 | 0.922 | 0.80<br>(89%) | Disease |
| - | Y328F | Disease | 2 | 0.592 | Disease | 7 | 0.829 | 0.62<br>(83%) | Disease |
| rs115923559<br>5 | W323R | Disease | 8 | 0.91 | Disease | 9 | 0.939 | 0.85<br>(91%) | Disease |
| rs128870697<br>2 | S298F | Disease | 6 | 0.787 | Disease | 7 | 0.866 | 0.86<br>(91%) | Disease |
| rs133959447<br>5 | G272R | Disease | 7 | 0.843 | Disease | 8 | 0.906 | 0.85<br>(91%) | Disease |
| rs139446877<br>0 | P248L | Disease | 4 | 0.711 | Disease | 2 | 0.589 | 0.80<br>(89%) | Disease |
| rs131263800<br>5 | V223E | Disease | 6 | 0.813 | Disease | 6 | 0.821 | 0.83<br>(90%) | Disease |
| rs749382362 | R219Q | Disease | 3 | 0.667 | Disease | 7 | 0.864 | 0.64<br>(84%) | Disease |
| rs374238430 | R219W | Disease | 5 | 0.773 | Disease | 8 | 0.909 | 0.67<br>(85%) | Disease |
| rs103983449<br>2 | V209E | Disease | 7 | 0.839 | Disease | 7 | 0.844 | 0.79<br>(89%) | Disease |
| rs747349523 | D203Y | Disease | 8 | 0.893 | Disease | 9 | 0.949 | 0.86<br>(91%) | Disease |
| rs776300430 | G202E | Disease | 6 | 0.783 | Disease | 7 | 0.843 | 0.66<br>(85%) | Disease |
| rs367552387 | R201Q | Disease | 5 | 0.766 | Disease | 6 | 0.792 | 0.80<br>(89%) | Disease |
| - | R201P | Disease | 7 | 0.865 | Disease | 8 | 0.883 | 0.86<br>(91%) | Disease |
| rs773117166 | R201W | Disease | 6 | 0.787 | Disease | 7 | 0.845 | 0.86<br>(91%) | Disease |
| rs565257577 | G162R | Disease | 6 | 0.823 | Disease | 7 | 0.866 | 0.84<br>(90%) | Disease |

In SNP&GO, PHD-SNP and P-Mut software’s were used to predict the association of SNPs with disease. According to SNP&GO, PHD-SNP and P-Mut (69, 69 and 69 SNPs respectively) were found to be disease related SNPs. We selected the triple disease related SNPs only in 3 softwares for further analysis by I-Mutant 3.0 ,(Table 3). While I-Mutant result revealed that the protein stability decreased (which destabilize the amino acid interaction) in the following SNPs: (R1137C, Y943N, I936T, A907T, Y898C, D886V, R749K ,R722Q, L716S,Q710R, R693W ,R689Q ,R689W, G576D ,P410L ,G396V ,L336S ,H329Q ,Y328S, S298F, G272R, P248L, V223E, R219Q, R219W, V209E, G202E, R201P ,R201Q ,R201W ,G162R). While (Y992C, Y943C, N888Y, Y797C, T793R, D791V, Y328C, Y328F, W323R, D203Y) are found to increase the protein stability (Table 3). Figures (1&2).

**Table 3:** stability analysis predicted by I-Mutant version 3.0 (Also show Most deleterious nsSNPs):

| dbSNP rs# | Amino Acid Change | SVM2 Prediction Effect | RI | DDG Value Prediction |
| --- | --- | --- | --- | --- |
| rs377679012 | R1137C | Decrease | 5 | -0.93 |
| rs372723869 | Y992C | Increase | 4 | -0.46 |
| rs924119518 | Y943C | Increase | 1 | -0.98 |
| rs1319436059 | Y943N | Decrease | 6 | -1.34 |
| rs199617267 | I936T | Decrease | 6 | -2.16 |
| rs1218321803 | A907T | Decrease | -0.7 | 6 |
| rs199679232 | Y898C | Decrease | -1.14 | 4 |
| rs752349406 | N888Y | Increase | 0.13 | 2 |
| rs1270126477 | D886V | Decrease | 0.04 | 4 |
| rs1181862976 | Y797C | Increase | -0.85 | 4 |
| rs201596987 | T793R | Increase | 0.02 | 4 |
| - | D791V | Increase | 0.63 | 2 |
| rs1452809051 | R749K | Decrease | -0.9 | 8 |
| rs370575901 | R722Q | Decrease | -0.88 | 9 |
| rs767003999 | L716S | Decrease | 9 | -2.22 |
| rs1275065654 | Q710R | Decrease | 1 | -0.2 |
| rs763692410 | R693W | Decrease | 5 | -0.48 |
| rs201742754 | R689Q | Decrease | 9 | -1.24 |
| rs201390288 | R689W | Decrease | 6 | -0.66 |
| rs56229130 | G576D | decrease | 6 | -1.02 |
| rs375957332 | P410L | decrease | 8 | -1.6 |
| rs1414061926 | G396V | decrease | 8 | -0.84 |
| rs775321513 | L336S | decrease | 8 | -1.89 |
| rs764033545 | H329Q | decrease | 3 | -0.28 |
| rs750663365 | Y328S | decrease | 7 | -0.41 |
| - | Y328C | Increase | 2 | -0.37 |
| - | Y328F | Increase | 3 | 0.34 |
| rs1159235595 | W323R | Increase | 8 | -1.11 |
| rs1288706972 | S298F | decrease | 2 | -0.13 |
| rs1339594475 | G272R | Decrease | -0.39 | 1 |
| rs1394468770 | P248L | Decrease | -0.49 | 5 |
| rs1312638005 | V223E | Decrease | -1.17 | 6 |
| rs749382362 | R219Q | Decrease | -1.16 | 9 |
| rs374238430 | R219W | Decrease | -0.55 | 6 |
| rs1039834492 | V209E | decrease | 8 | -1.18 |
| rs747349523 | D203Y | increase | 0 | -0.17 |
| rs776300430 | G202E | decrease | 2 | -0.51 |
| rs367552387 | R201P | decrease | 6 | -0.72 |
| - | R201Q | decrease | 8 | -0.93 |
| rs773117166 | R201W | decrease | 4 | -0.22 |
| rs565257577 | G162R | decrease | 3 | -0.36 |

GeneMANIA revealed that *IKBKAP* gene has many important functions : DNA-directed RNA polymerase complex, DNA-directed RNA polymerase II, holoenzyme, DNA-templated transcription, elongation, nuclear DNA-directed RNA polymerase complex, positive regulation of cell migration, positive regulation of cell motility, positive regulation of locomotion, RNA polymerase complex, transcription elongation factor complex, transcription elongation from RNA polymerase II promoter. The genes co-expressed with, share similar protein domain, or participate to achieve similar function are illustrated by GeneMANIA and shown in figures (4 to 44) Table (4 & 5).

**Table 4:** The *IKBKAP* gene functions and its appearance in network and genome:

| Function | FDR | Genes in network | Genes in genome |
| --- | --- | --- | --- |
| transcription elongation factor complex | 0.000385 | 4 | 28 |
| transcription elongation from RNA polymerase II promoter | 0.009942 | 4 | 75 |
| DNA-directed RNA polymerase II, holoenzyme | 0.009942 | 4 | 81 |
| nuclear DNA-directed RNA polymerase complex | 0.009942 | 4 | 95 |
| RNA polymerase complex | 0.009942 | 4 | 96 |
| DNA-directed RNA polymerase complex | 0.009942 | 4 | 95 |
| DNA-templated transcription, elongation | 0.013634 | 4 | 108 |
| positive regulation of locomotion | 0.125453 | 4 | 213 |
| positive regulation of cell motility | 0.125453 | 4 | 204 |
| positive regulation of cell migration | 0.125453 | 4 | 199 |

**\*FDR:** false discovery rate is greater than or equal to the probability that this is a false positive.

**Table 5:** The gene co-expressed, share domain and Interaction with *IKBKAP* gene network:(Also show The 41 *novel mutations*):

| Gene 1 | Gene 2 | Weight | Network group |
| --- | --- | --- | --- |
| ELP2 | ELP1 | 0.009 | Co-expression |
| GAPVD1 | ELP1 | 0.010505 | Co-expression |
| URM1 | PPP6C | 0.012673 | Co-expression |
| GAPVD1 | ELP1 | 0.019938 | Co-expression |
| CD2AP | ELP1 | 0.010033 | Co-expression |
| URM1 | PPP6C | 0.014834 | Co-expression |
| GAPVD1 | ELP1 | 0.020476 | Co-expression |
| GAPVD1 | PPP6C | 0.017376 | Co-expression |
| FOCAD | ELP1 | 0.010588 | Co-expression |
| IKBKB | ELP1 | 0.009126 | Co-expression |
| ELP5 | ELP6 | 0.017894 | Co-expression |
| IRF4 | NFKBIA | 0.017726 | Co-expression |
| ELP2 | ELP1 | 0.006755 | Co-expression |
| DMAP1 | ELP1 | 0.013834 | Co-expression |
| ELP4 | PPP6C | 0.009147 | Co-expression |
| IRF4 | NFKBIA | 0.013234 | Co-expression |
| NFKBIA | ELP1 | 0.315041 | Pathway |
| IKBKB | NFKBIA | 0.012293 | Pathway |
| MAP3K14 | IKBKB | 0.013495 | Pathway |
| SHC1 | ELP1 | 0.126671 | Physical Interactions |
| ELP3 | ELP2 | 0.181392 | Physical Interactions |
| TSC22D1 | ELP1 | 0.331631 | Physical Interactions |
| NUP50 | ELP1 | 0.277373 | Physical Interactions |
| NUP50 | TSC22D1 | 0.157924 | Physical Interactions |
| CD2AP | ELP1 | 0.137789 | Physical Interactions |
| CD2AP | ELP2 | 0.049176 | Physical Interactions |
| CD2AP | TSC22D1 | 0.078451 | Physical Interactions |
| ELP2 | ELP1 | 0.333333 | Physical Interactions |
| ELP3 | ELP1 | 0.333333 | Physical Interactions |
| ELP3 | ELP2 | 0.333333 | Physical Interactions |
| ELP4 | ELP1 | 0.333333 | Physical Interactions |
| ELP4 | ELP2 | 0.333333 | Physical Interactions |
| ELP4 | ELP3 | 0.333333 | Physical Interactions |
| NFKBIA | ELP1 | 0.215941 | Physical Interactions |
| IKBKB | NFKBIA | 0.037527 | Physical Interactions |
| URM1 | ELP1 | 0.141754 | Physical Interactions |
| MAP3K14 | IKBKB | 0.048823 | Physical Interactions |
| IRF4 | ELP1 | 0.407145 | Physical Interactions |
| ELP2 | ELP1 | 0.133207 | Physical Interactions |
| ELP3 | ELP1 | 0.231538 | Physical Interactions |
| ELP3 | ELP2 | 0.344877 | Physical Interactions |
| ELP4 | ELP2 | 0.48935 | Physical Interactions |
| DPH3 | ELP1 | 0.517968 | Physical Interactions |
| FOCAD | ELP1 | 0.517968 | Physical Interactions |
| ELP2 | ELP1 | 0.0749 | Physical Interactions |
| ELP3 | ELP1 | 0.083596 | Physical Interactions |
| ELP3 | ELP2 | 0.1826 | Physical Interactions |
| NFKBIA | ELP1 | 0.014861 | Physical Interactions |
| ELP4 | ELP1 | 0.104887 | Physical Interactions |
| ELP4 | ELP2 | 0.229106 | Physical Interactions |
| ELP4 | ELP3 | 0.255706 | Physical Interactions |
| IKBKB | ELP1 | 0.011662 | Physical Interactions |
| IKBKB | NFKBIA | 0.005054 | Physical Interactions |
| ELP6 | ELP1 | 0.286862 | Physical Interactions |
| MAP3K14 | ELP1 | 0.038893 | Physical Interactions |
| MAP3K14 | NFKBIA | 0.016857 | Physical Interactions |
| MAP3K14 | IKBKB | 0.013228 | Physical Interactions |
| ELP5 | ELP1 | 0.170383 | Physical Interactions |
| IKBKB | ELP1 | 0.255299 | Physical Interactions |
| IKBKB | NFKBIA | 0.058324 | Physical Interactions |
| MAP3K14 | IKBKB | 0.643594 | Physical Interactions |
| ELP6 | ELP4 | 0.312748 | Physical Interactions |
| ELP5 | ELP6 | 0.312748 | Physical Interactions |
| ELP3 | ELP1 | 0.252782 | Physical Interactions |
| NFKBIA | ELP1 | 0.016702 | Physical Interactions |
| NFKBIA | ELP3 | 0.139754 | Physical Interactions |
| IRF4 | ELP1 | 0.0903 | Physical Interactions |
| IKBKB | NFKBIA | 0.011146 | Physical Interactions |
| MAP3K14 | IKBKB | 0.006473 | Physical Interactions |
| ELP5 | ELP4 | 0.751237 | Physical Interactions |
| ELP5 | ELP6 | 0.583434 | Physical Interactions |
| IKBKB | ELP1 | 0.022347 | Physical Interactions |
| MAP3K14 | ELP1 | 0.025064 | Physical Interactions |
| MAP3K14 | IKBKB | 0.01664 | Physical Interactions |
| ELP2 | ELP1 | 0.5 | Predicted |
| ELP3 | ELP1 | 0.5 | Predicted |
| ELP3 | ELP2 | 0.5 | Predicted |
| ELP2 | ELP1 | 0.108428 | Predicted |
| ELP3 | ELP1 | 0.095014 | Predicted |
| ELP3 | ELP2 | 0.106134 | Predicted |
| PPP6C | ELP1 | 0.040139 | Predicted |
| DPH3 | ELP1 | 0.092118 | Predicted |
| DPH3 | ELP2 | 0.1029 | Predicted |
| DPH3 | ELP3 | 0.090169 | Predicted |
| KTI12 | ELP1 | 0.161683 | Predicted |
| KTI12 | ELP2 | 0.180607 | Predicted |
| KTI12 | ELP3 | 0.158263 | Predicted |
| DMAP1 | ELP1 | 0.092452 | Predicted |
| PPP6C | ELP1 | 1 | Predicted |
| DPH3 | ELP2 | 0.707107 | Predicted |
| DPH3 | ELP3 | 0.707107 | Predicted |
| ELP2 | ELP1 | 0.53965 | Predicted |
| ELP4 | ELP1 | 0.53965 | Predicted |
| IKBKB | ELP1 | 0.081517 | Predicted |
| MAP3K14 | ELP1 | 0.126572 | Predicted |
| MAP3K14 | NFKBIA | 0.068474 | Predicted |
| ELP2 | ELP1 | 0.393402 | Predicted |
| ELP3 | ELP1 | 0.451851 | Predicted |
| ELP3 | ELP2 | 0.290279 | Predicted |
| KTI12 | ELP2 | 0.66162 | Predicted |
| MAP3K14 | IKBKB | 0.00439 | Shared protein domains |

**Figure 4:**
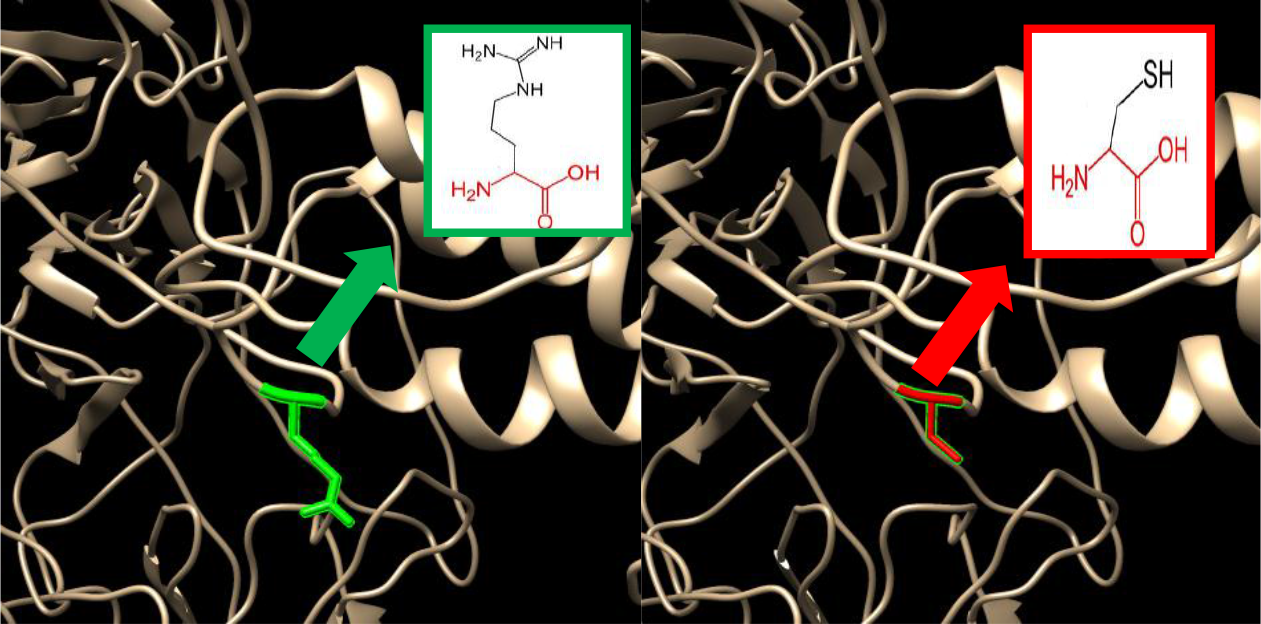
(rs377679012) :(R1137C): The amino acid Arginine change to Cysteine at position 1137.

**Figure 5:**
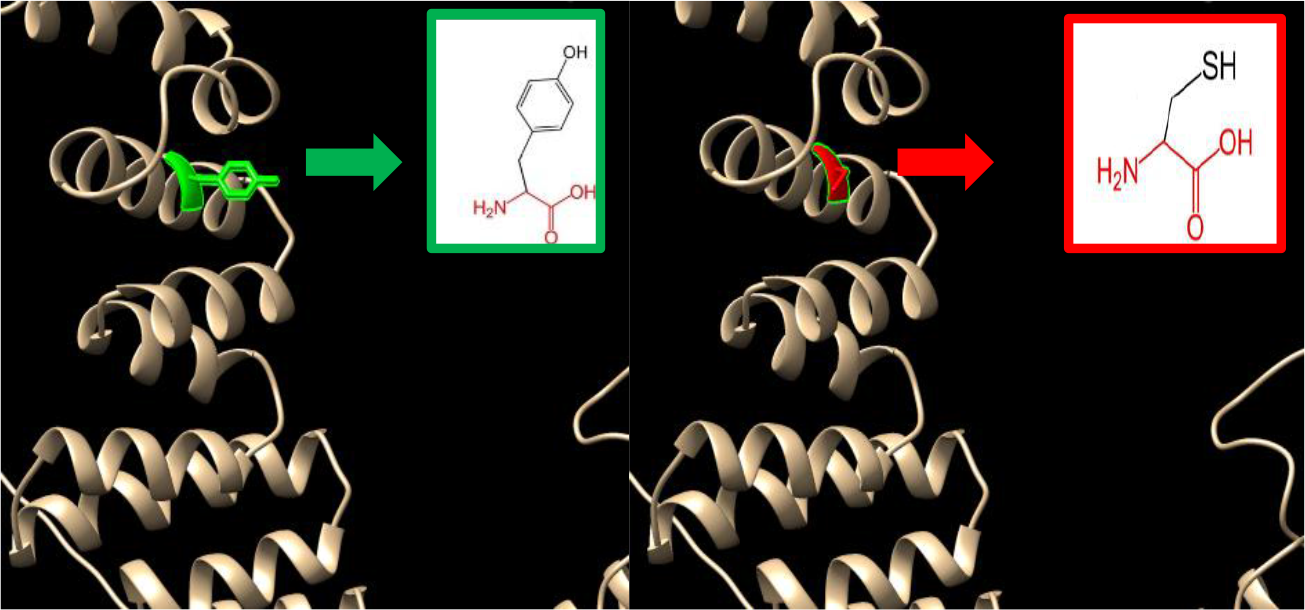
(rs372723869) :(Y992C): The amino acid Tyrosine change to Cysteine at position 992.

**Figure 6:**
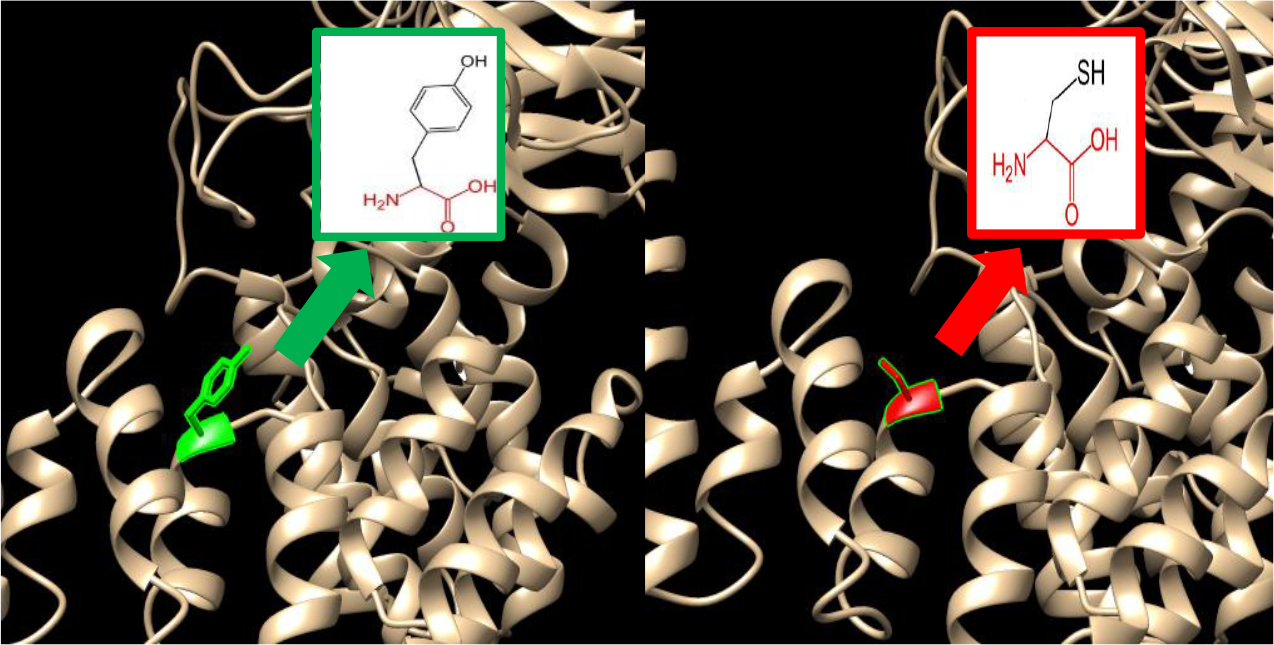
(rs924119518) :(Y943C): The amino acid Tyrosine change to Cysteine at position 943.

**Figure 7:**
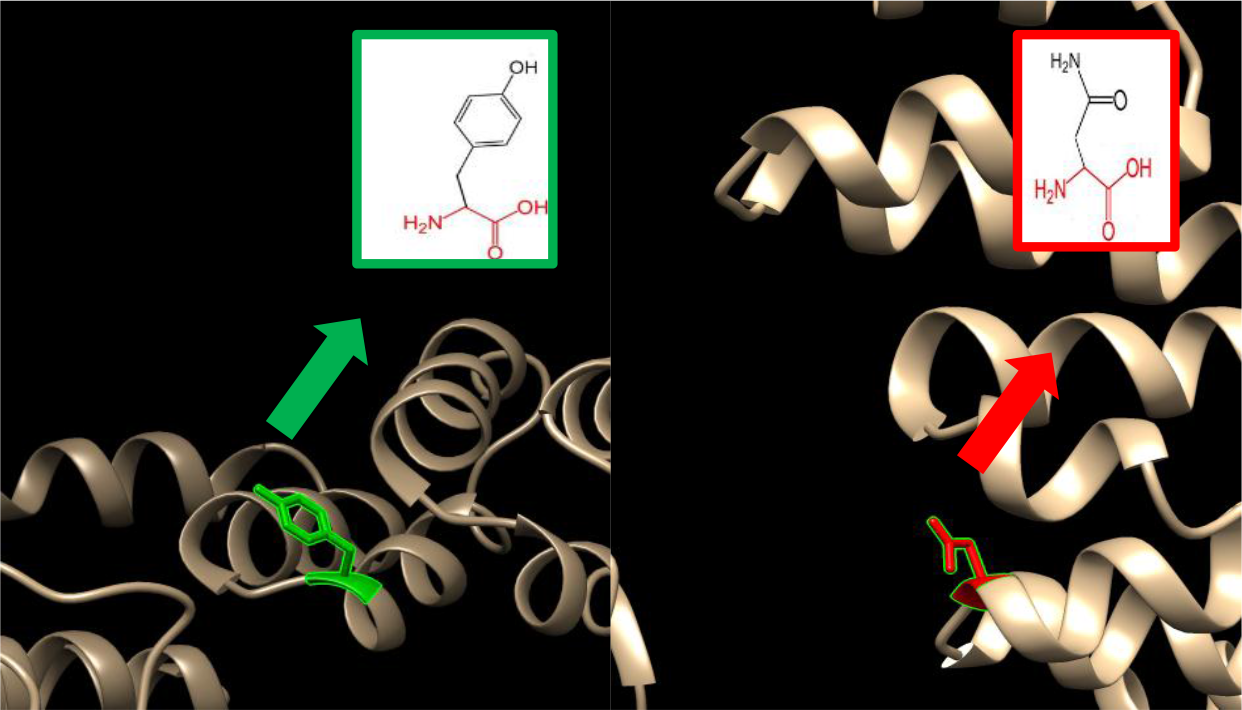
(rs1319436059) :(Y943N): The amino acid Tyrosine change to Asparagine at position 943.

**Figure 8::**
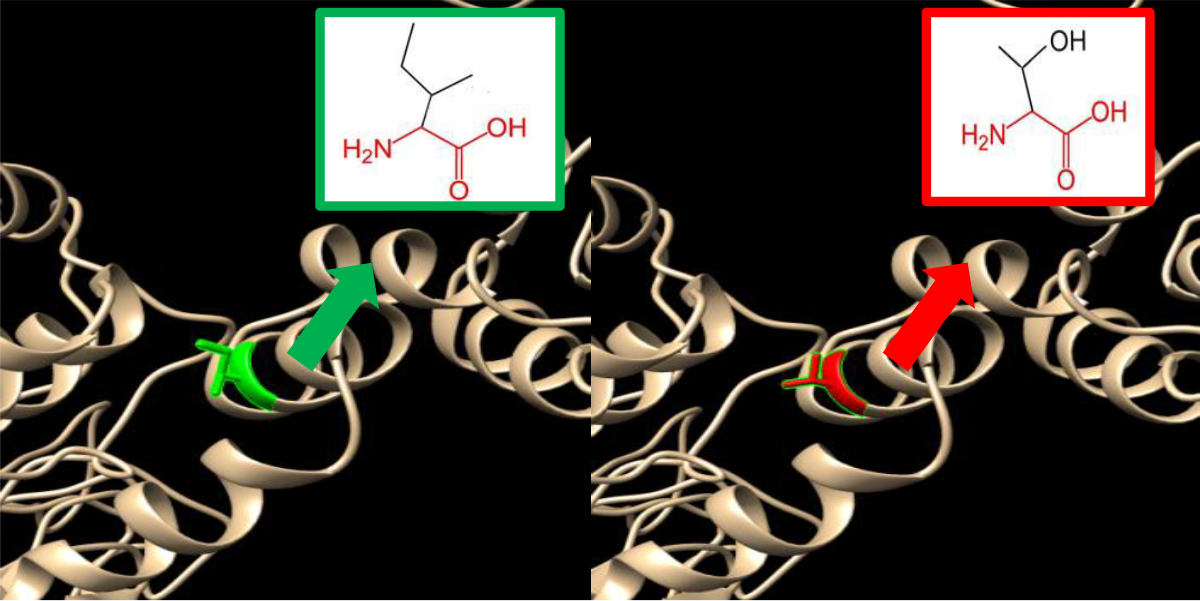
(rs199617267) :(I936T): The amino acid Isoleucine change to Threonine at position 936.

**Figure 9:**
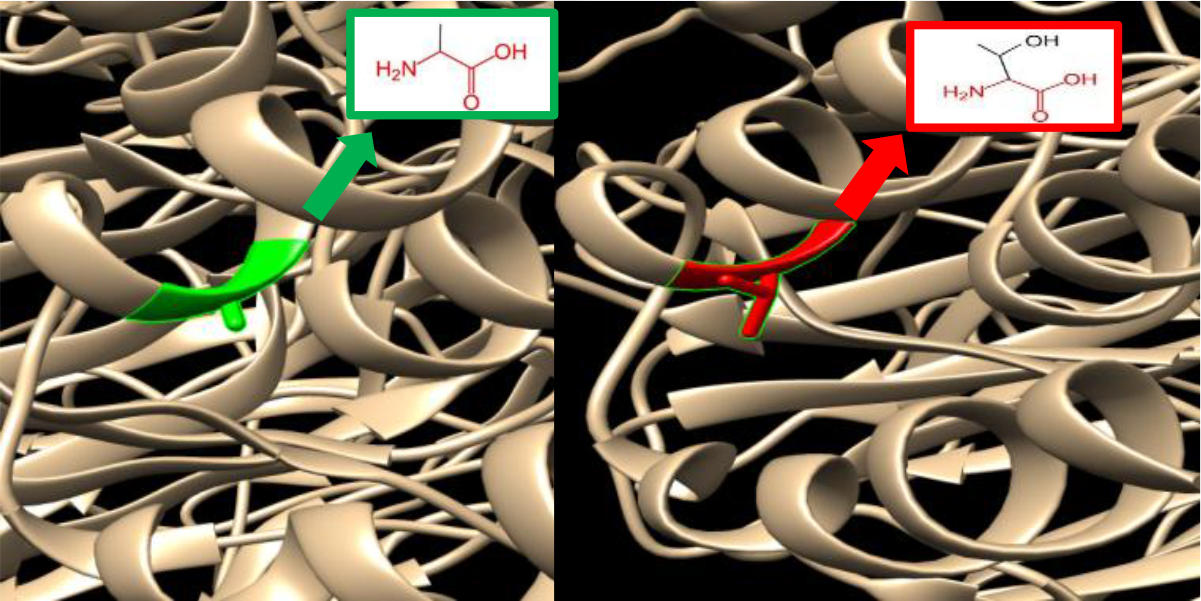
(rs1218321803): (A907T): The amino acid alanine to threonine at postion907.

**Figure 10:**
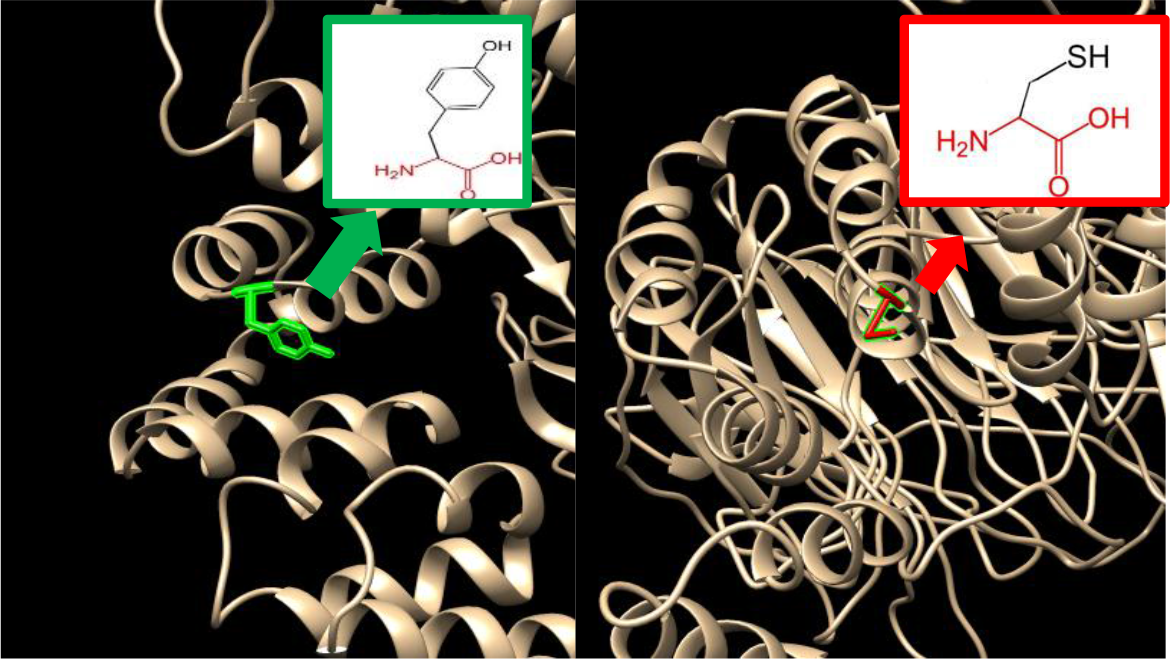
(rs199679232): (Y898C): The amino acid Tyrosine change to Cysteine at position 898.

**Figure 11:**
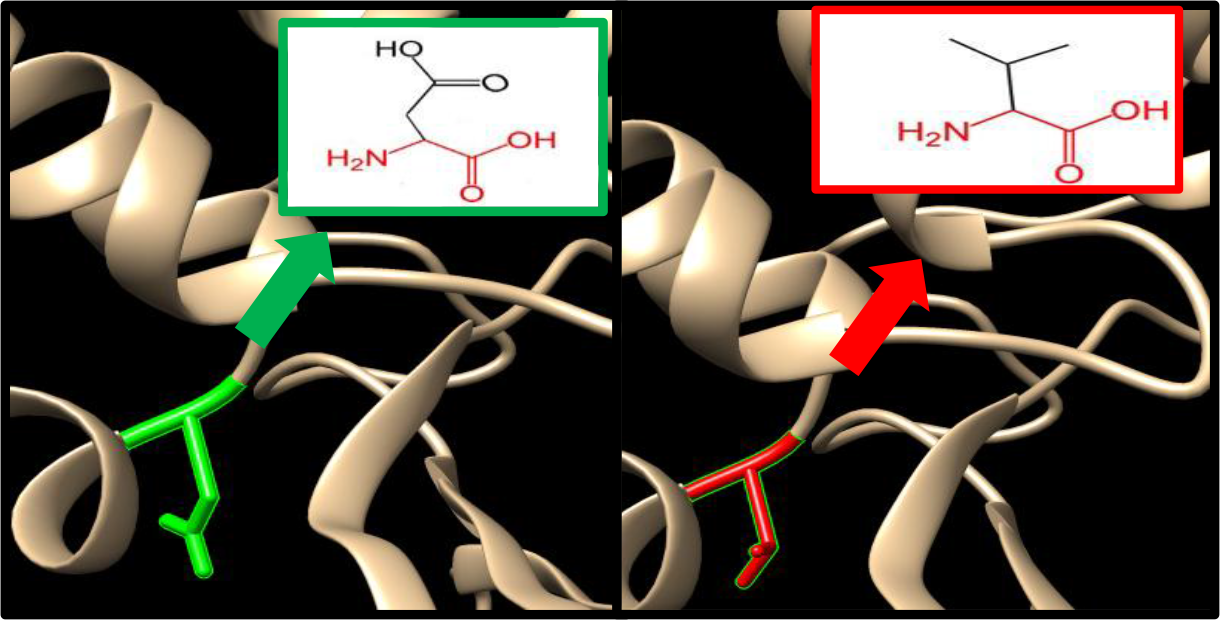
(rs752349406):(N888Y)The amino acid Asparagine change toTyrosine at position 888.

**Figure 12:**
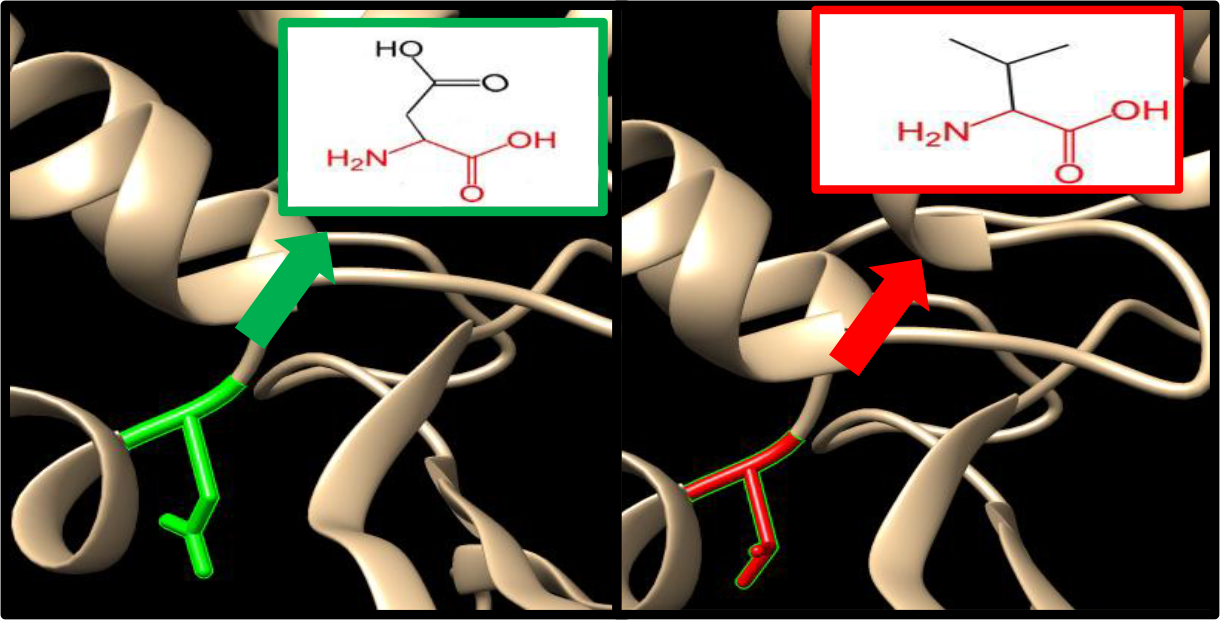
(rs1270126477):(D886V) The amino acid Aspartic acid change to Valine at position 886.

**Figure 13:**
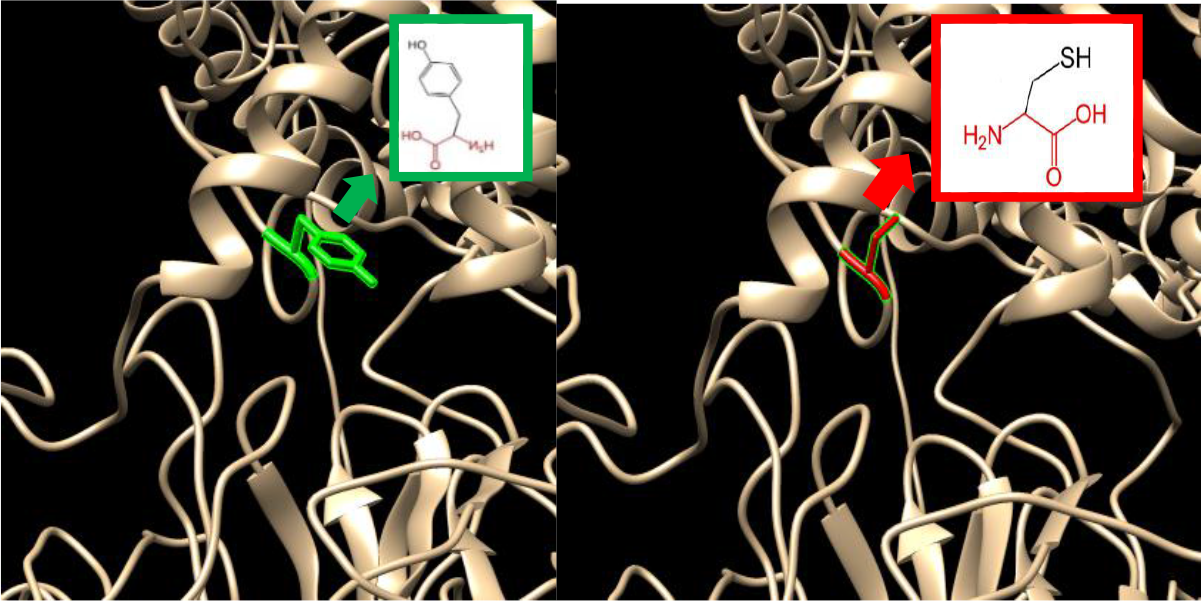
(rs1181862976) :(Y797C): The amino acid Tyrosine change to Cysteine at position797.

**Figure 14:**
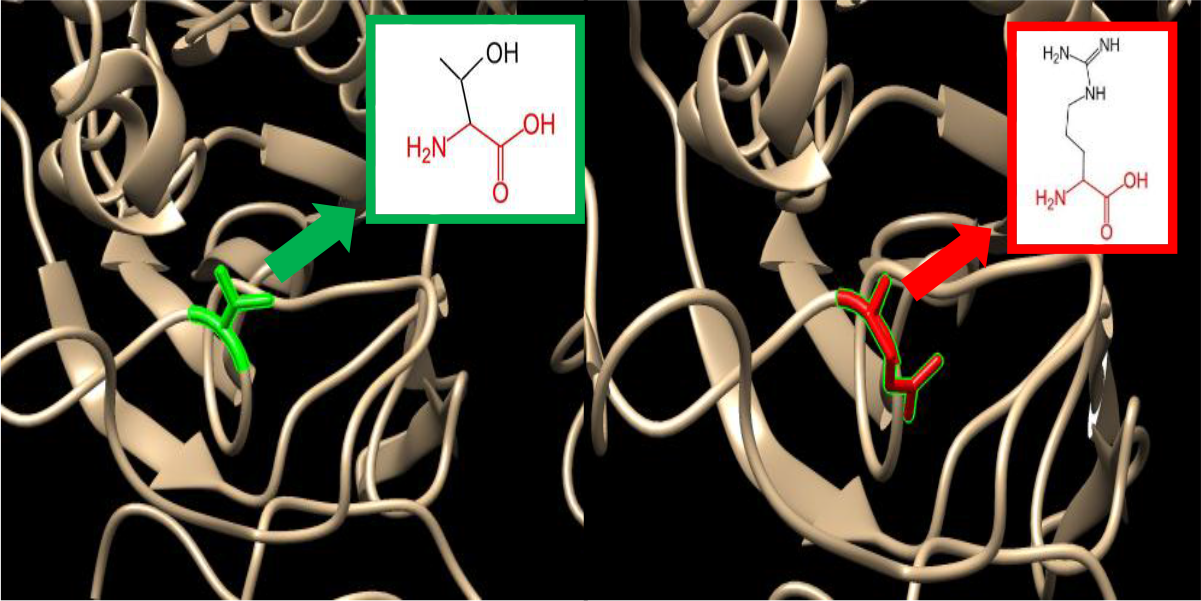
(rs201596987) :(T793R): The amino acid Threonine change to Arginine at position 793.

**Figure 15:**
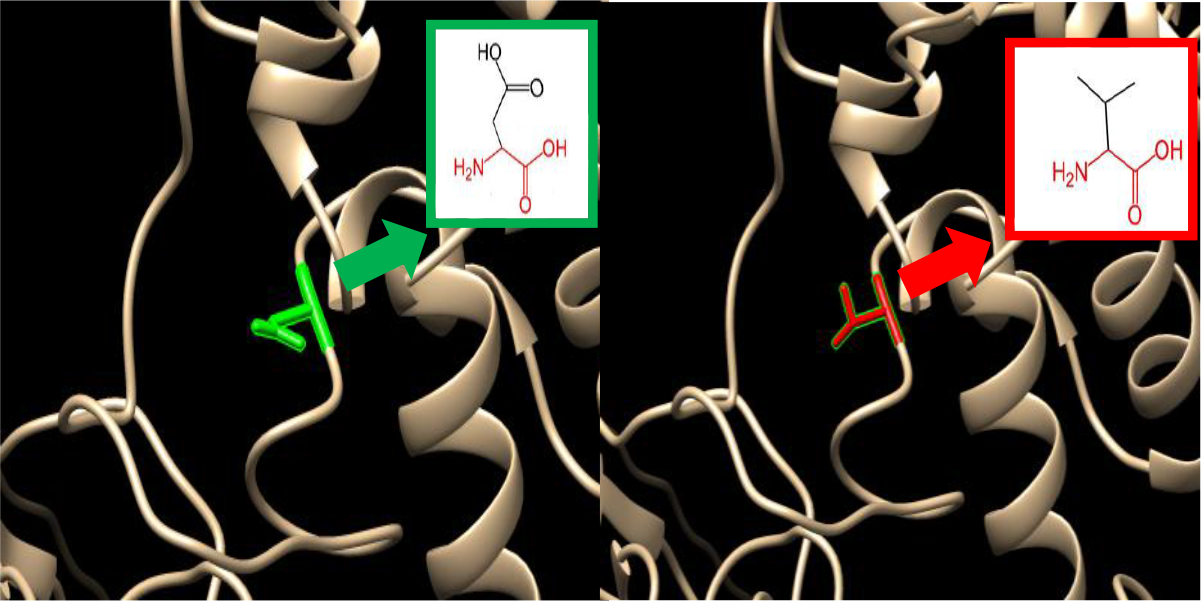
(D791V): The amino acid Aspartic acid change to Valine at position 791.

**Figure 16:**
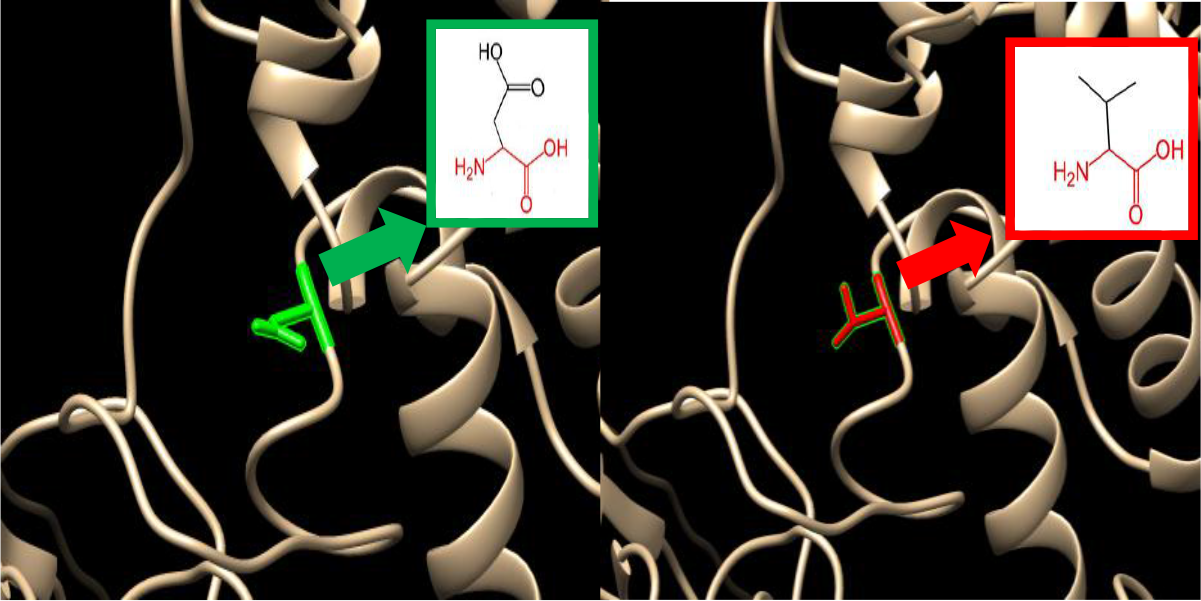
((rs1452809051):(R749K): The amino acid Arginine change to Lysine at position 749.

**Figure 17:**
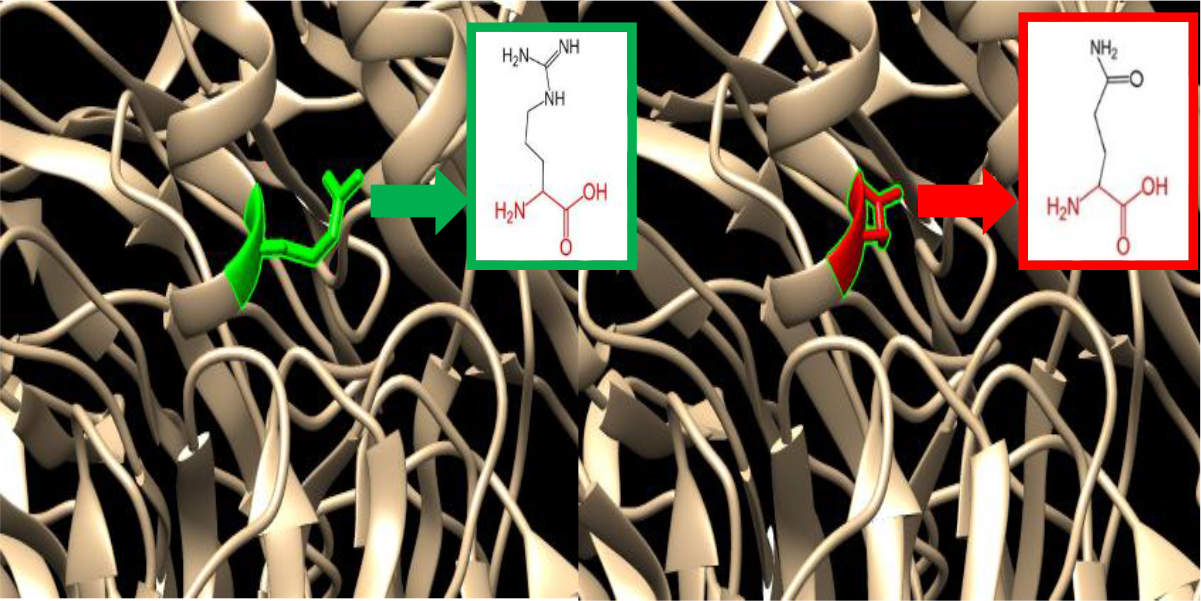
(rs1452809051) (R722Q): The amino acid Arginine change to Glutamine at position 722.

**Figure 18:**
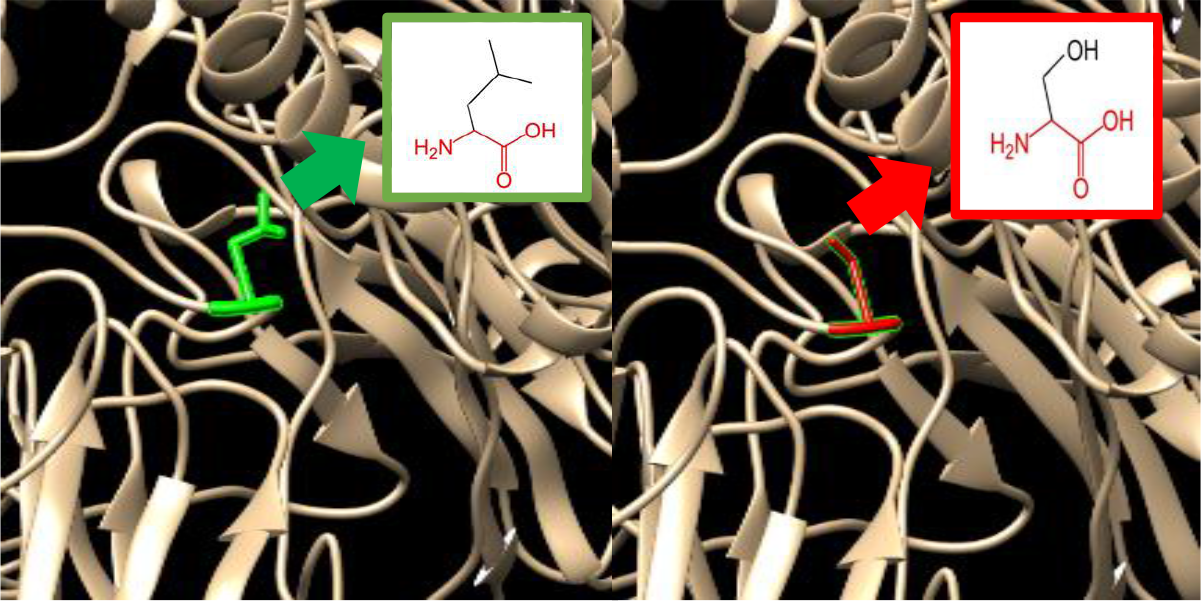
(rs767003999): (L716S): The amino acid Leucine change to serine at position 716.

**Figure 19:**
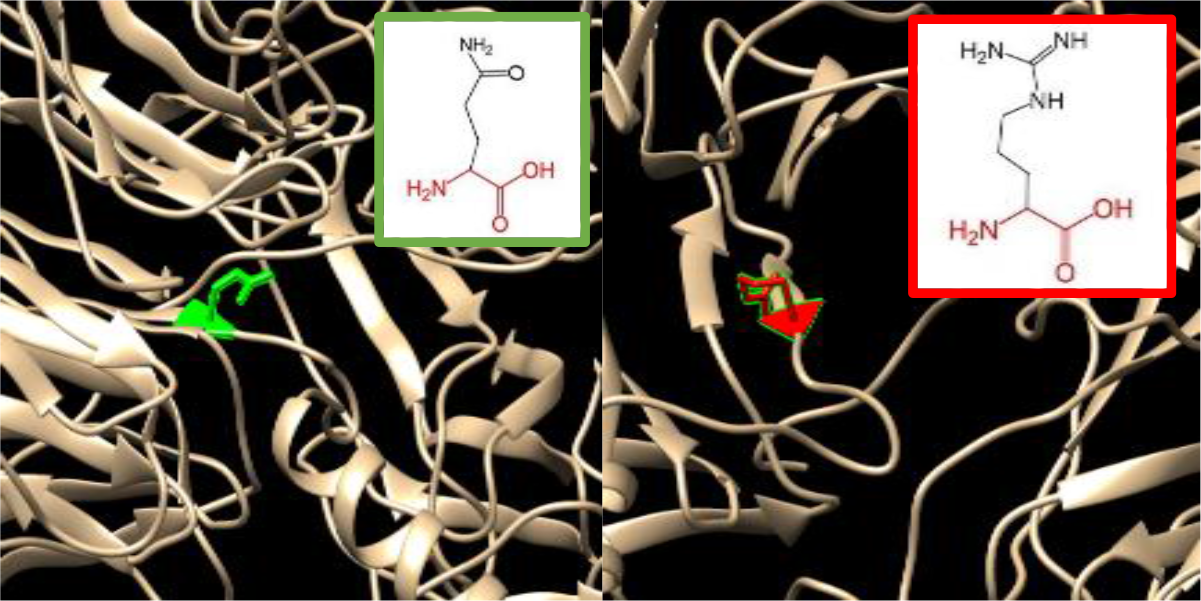
(rs1275065654): (Q710R): The amino acid Glutamine change to Arginine at position 710.

**Figure 20:**
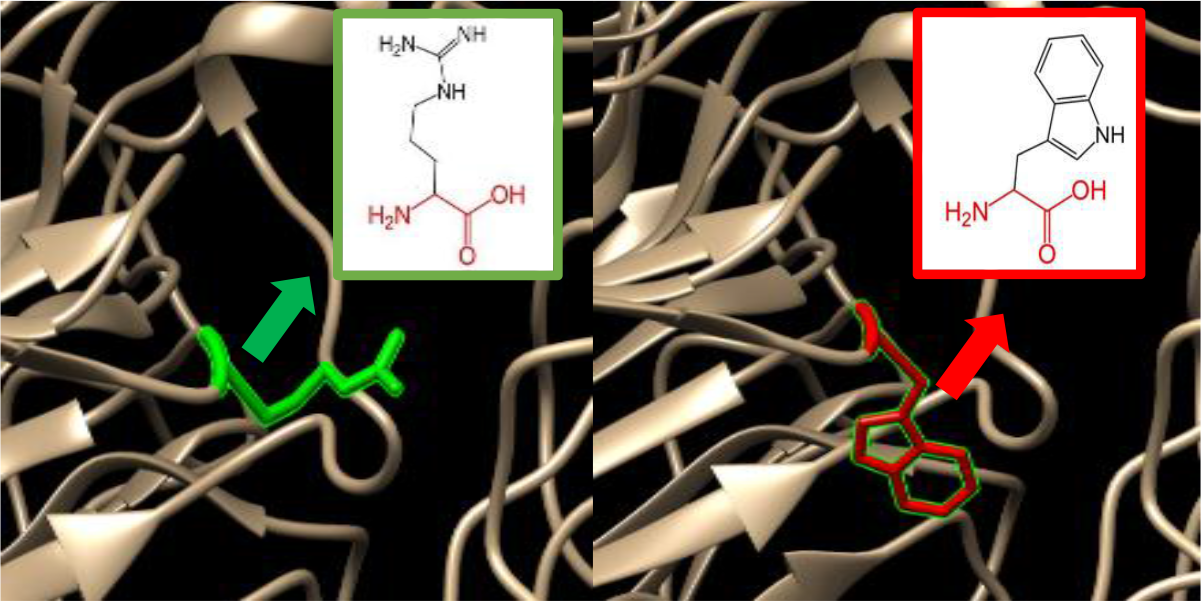
(rs763692410) :(R693W): The amino acid arginine change to Tryptophan at position 693.

**Figure 21:**
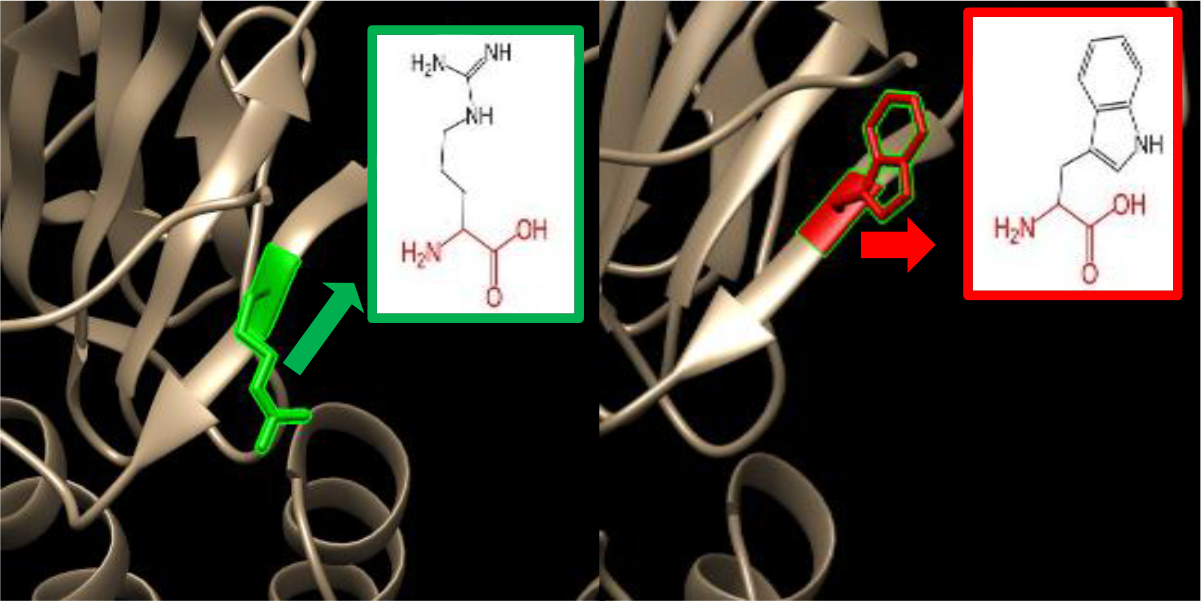
(rs201742754): (R689Q): The amino acid Arginine change to Glutamine at position 689.

**Figure 22:**
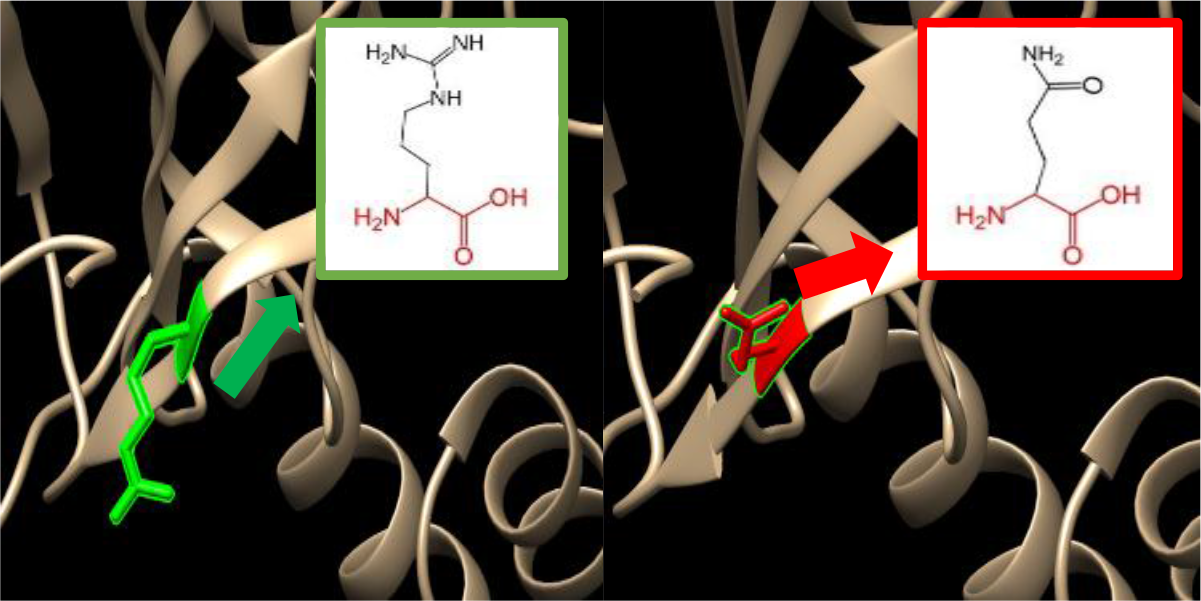
(rs201390288): (R689W): The amino acids Arginine change to Tryptophan at position 689.

**Figure 23:**
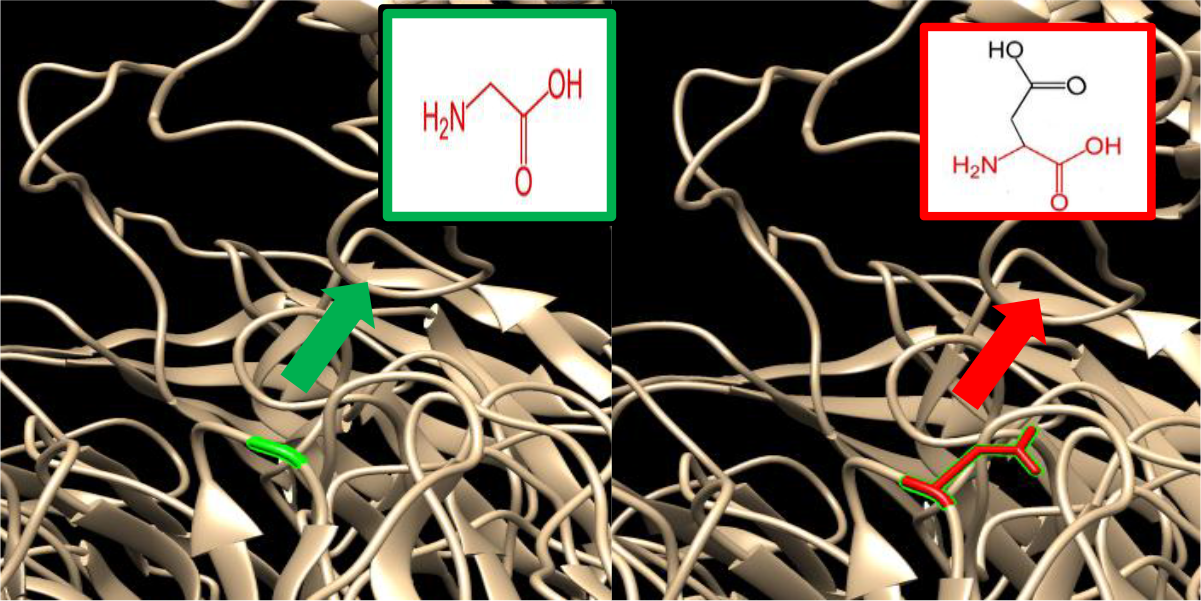
(rs56229130): (G576D): The amino acid Glycine changed to Aspartic acid at position 576.

**Figure 24:**
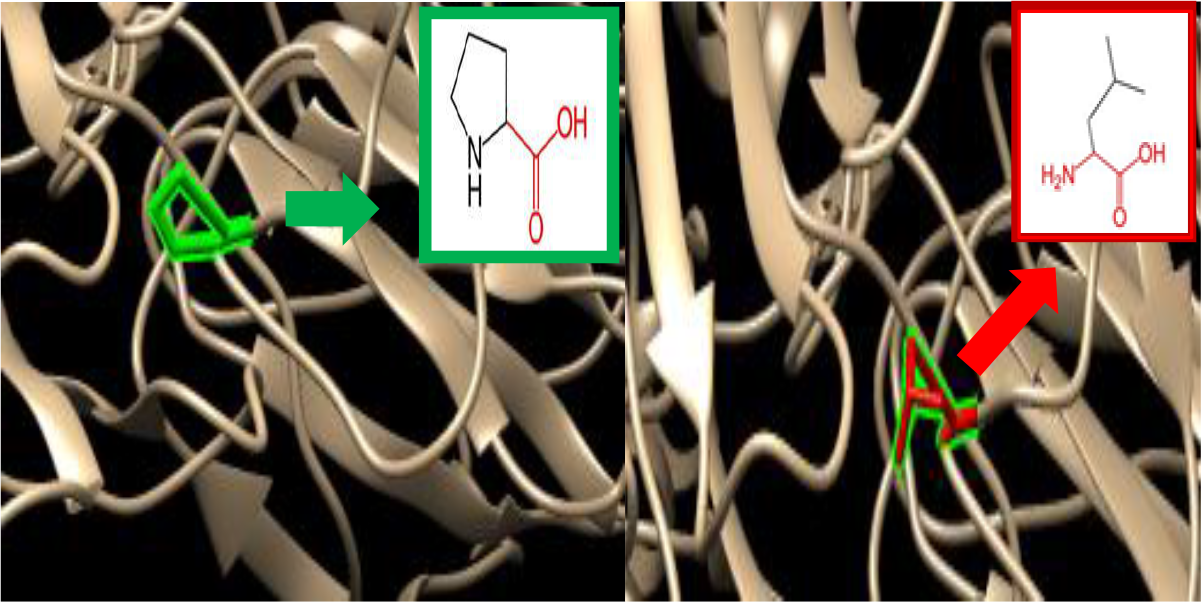
(rs375957332):(P410L) : The amino acid Proline changed to Leucine at position 410 .

**Figure 25:**
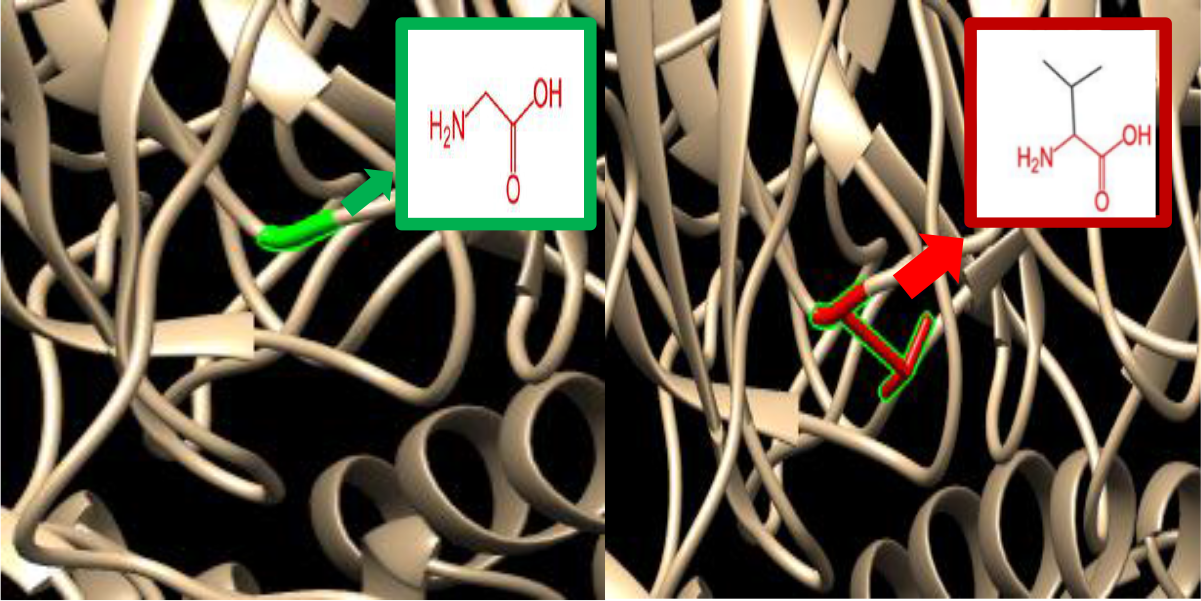
(rs1414061926): (G396V): The amino acid Glycine changes to Valine at position 396.

**Figure 26:**
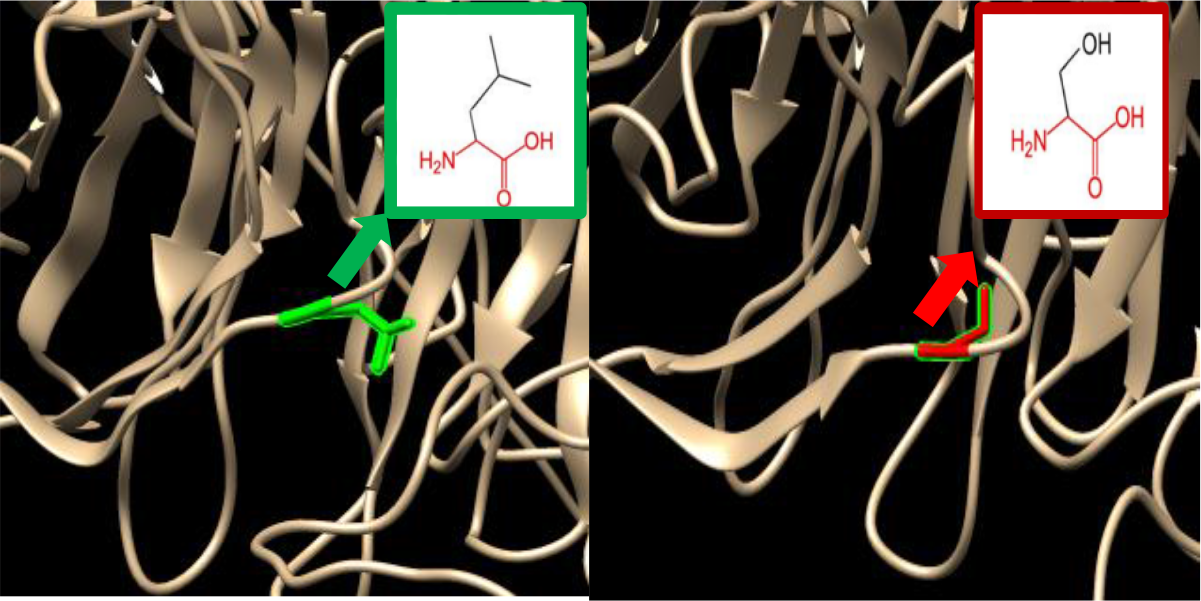
(rs775321513): (L336S): The amino acid Leucine changes to serine at position 336.

**Figure 27:**
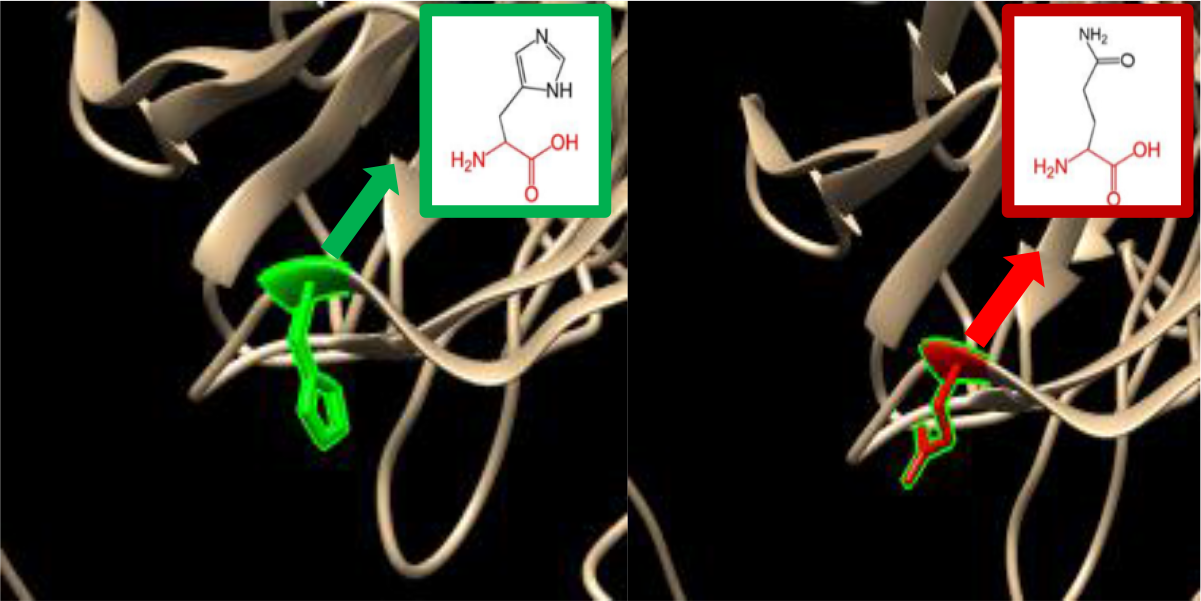
(rs764033545): (H329Q): The amino acids Histidine changes to glutamine at position 329.

**Figure 28:**
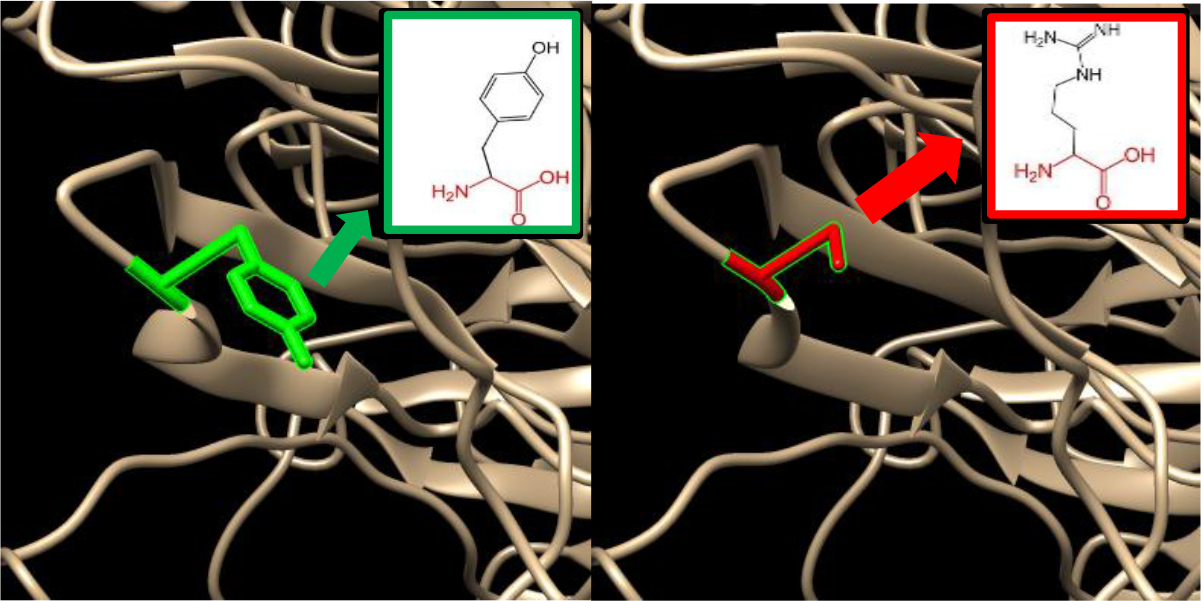
(rs750663365): (Y328S): The amino acid Tyrosine change to Serine at position 328.

**Figure 29:**
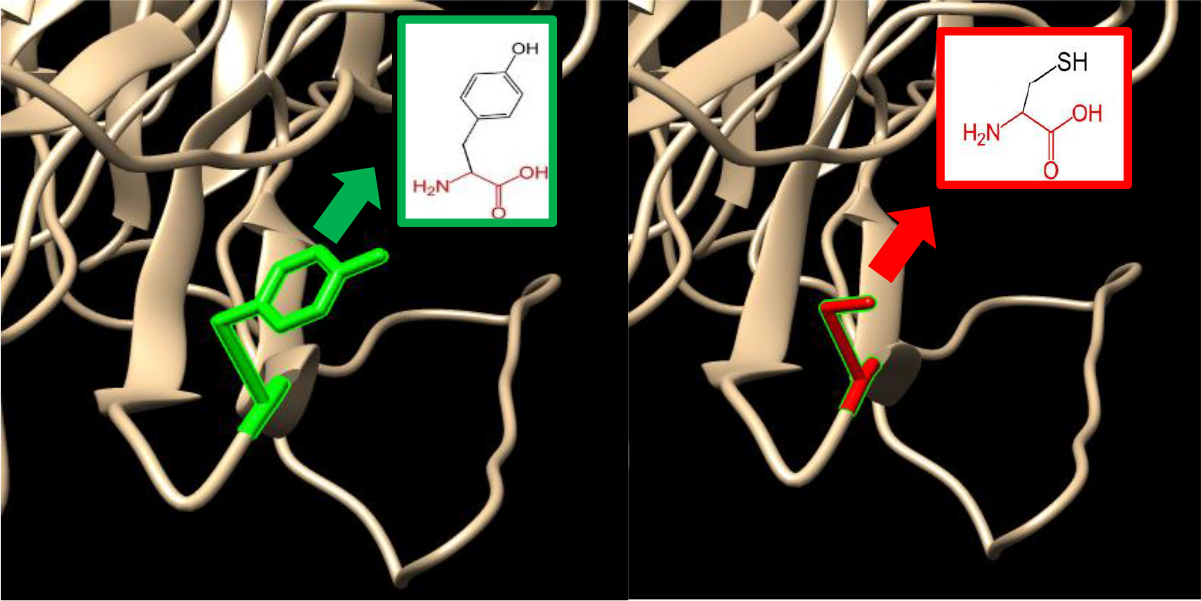
(Y328C): The amino acid Tyrosine change to Cysteine at position 328.

**Figure 30:**
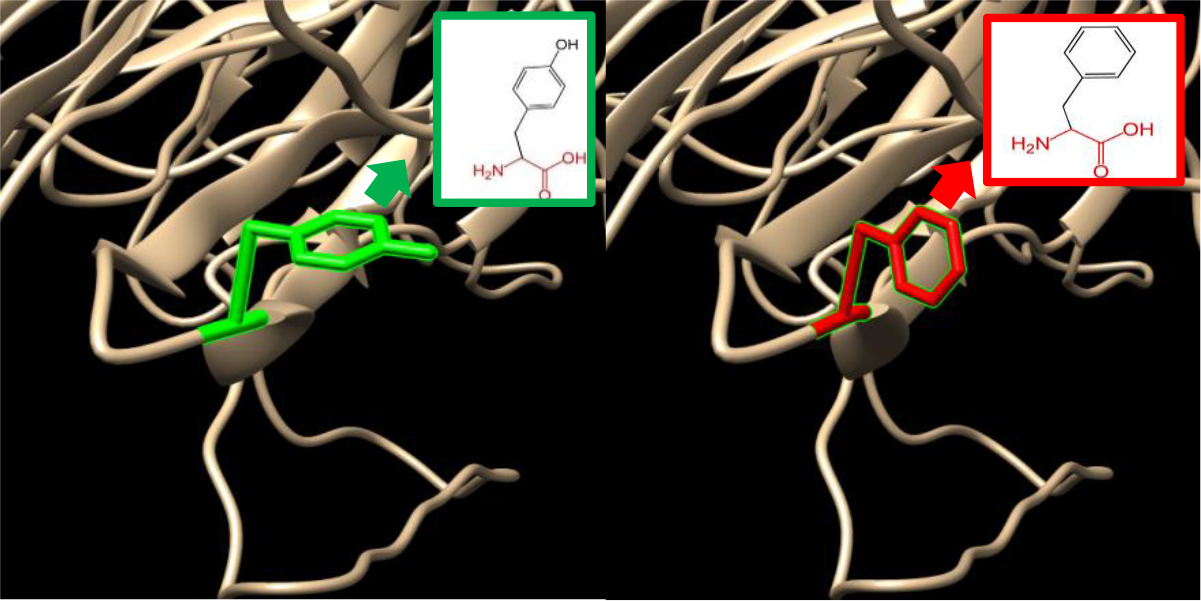
(Y328F): The amino acid Tyrosine changes to Phenylalanine at position 328.

**Figure 31:**
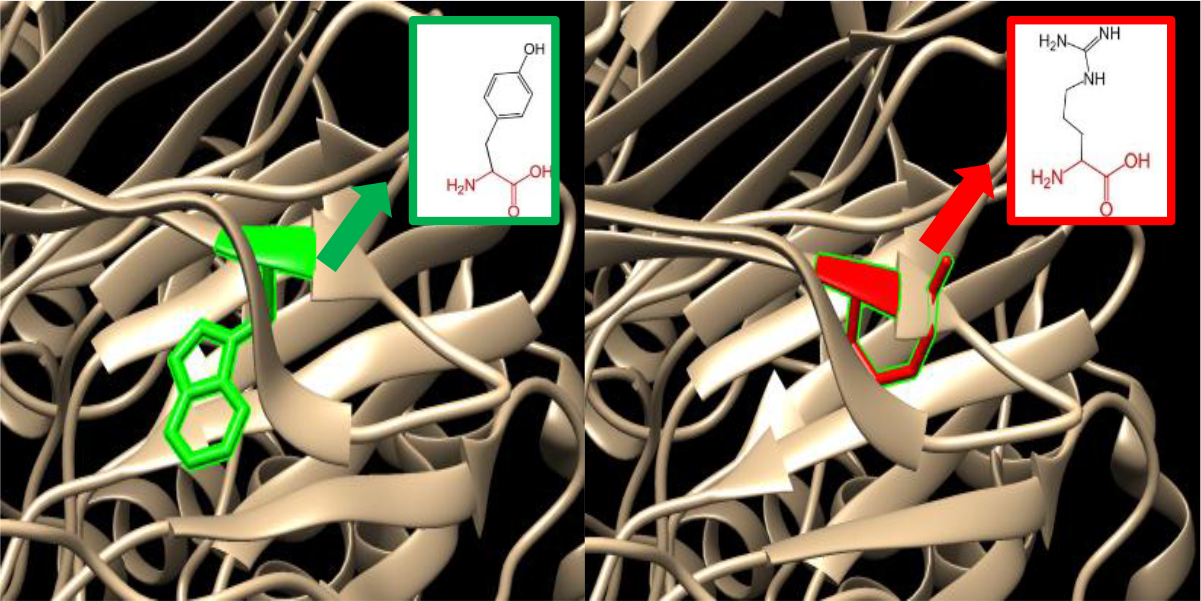
(rs1159235595): (W323R): The amino acid Tryptophan changes to Arginine at position 323.

**Figure 32:**
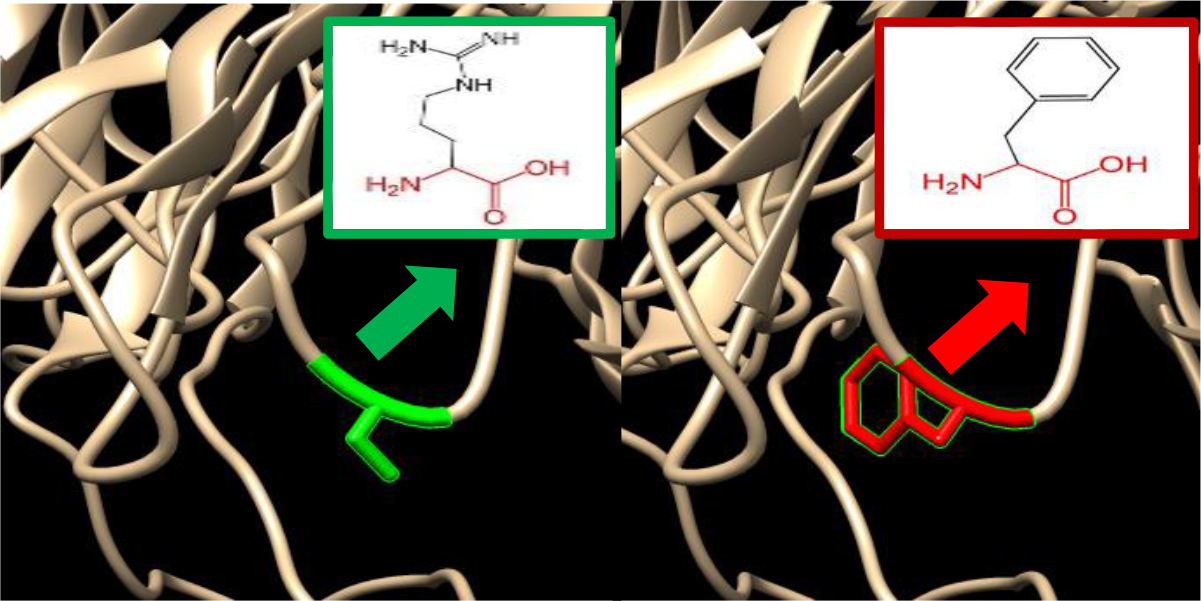
(rs1288706972): (S298F): The amino acid serine change to Phenylalanine at position 298.

**Figure 33:**
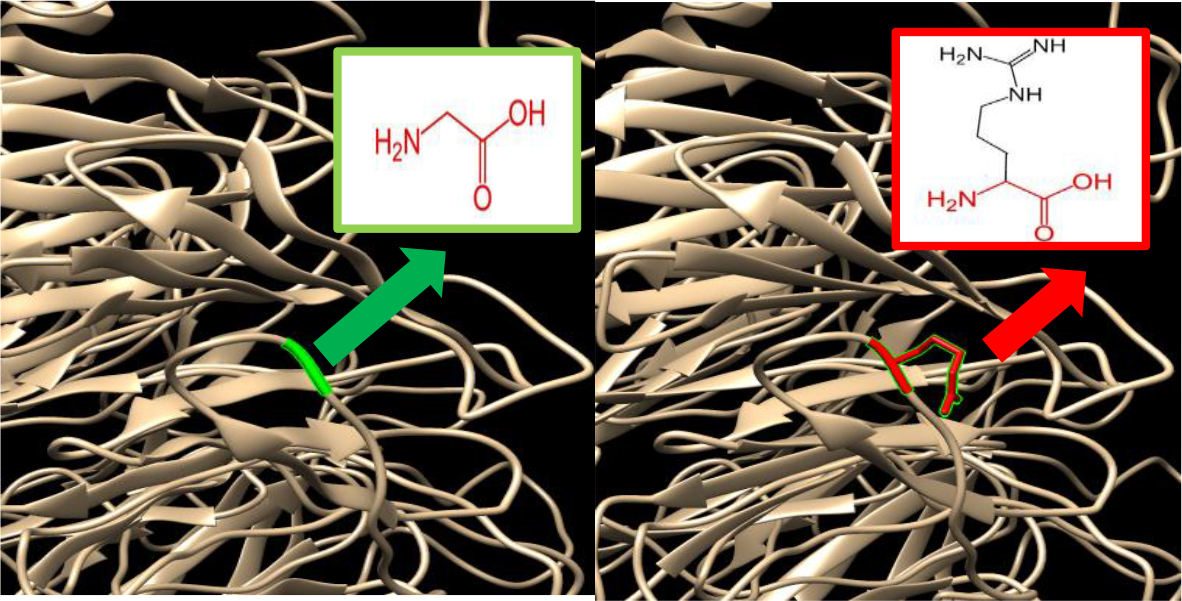
(rs1339594475): (G272R): The amino acid Glycine change to Arginine at position 272.

**Figure 34:**
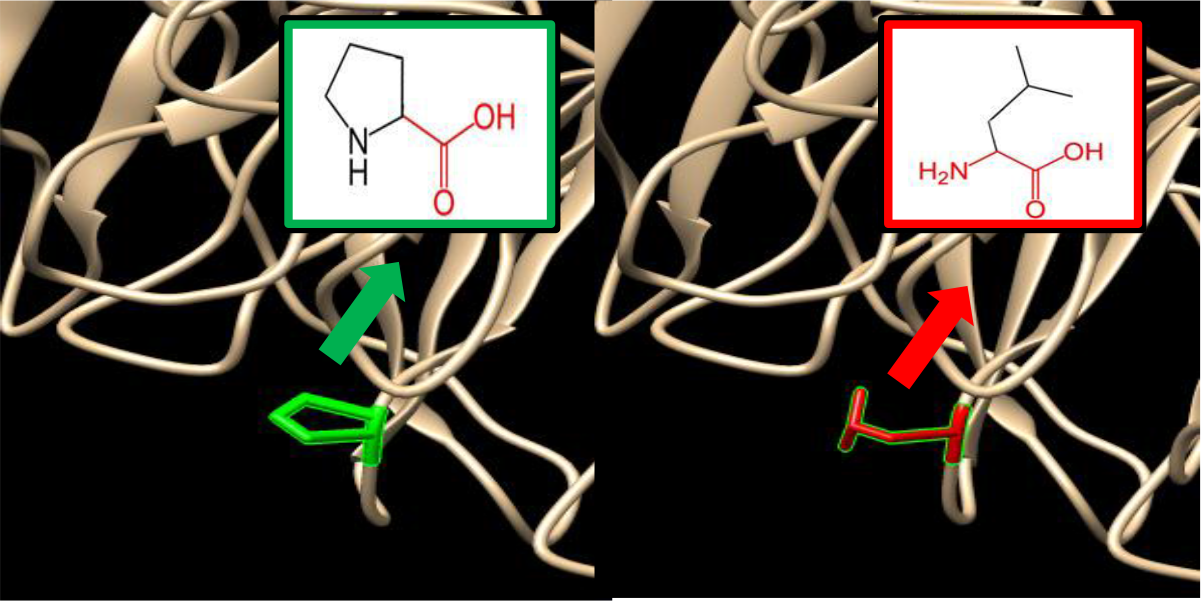
(rs1394468770): (P248L): The amino acid Proline changes to Leucine at position 248.

**Figure 35:**
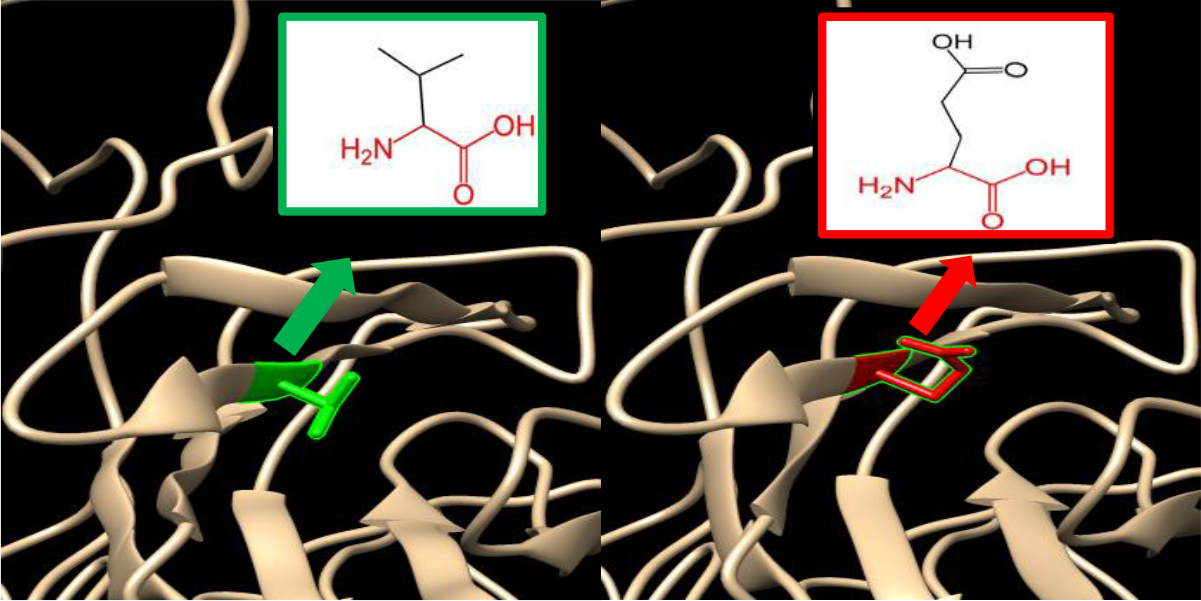
(rs1312638005): (V223E): The amino acid Valine change to Glutamic acid at position 223.

**Figure 36:**
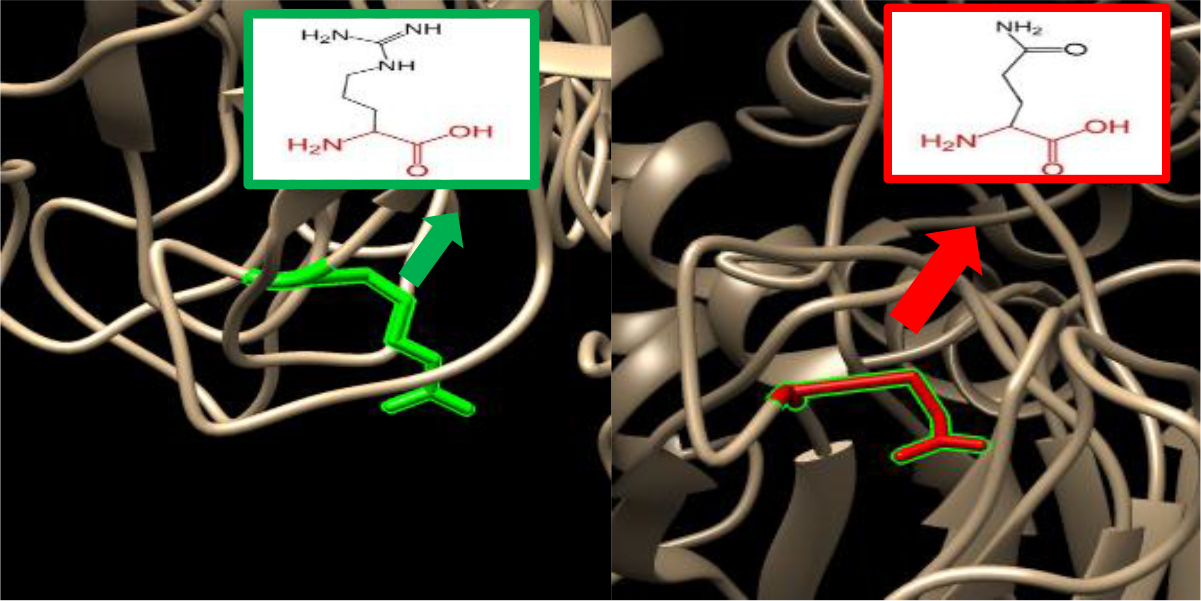
(rs749382362): (R219Q): The amino acid Arginine change to Glutamine at position 219.

**Figure 37:**
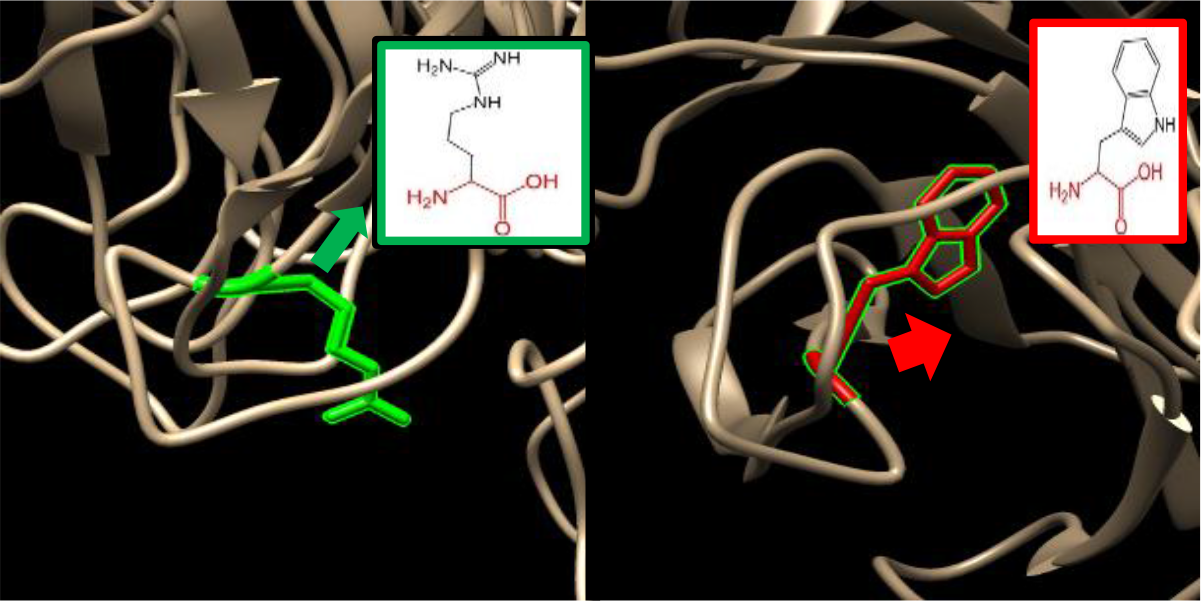
(rs374238430): (R219W): The amino acid Arginine changes to Tryptophan at position 219.

**Figure 38:**
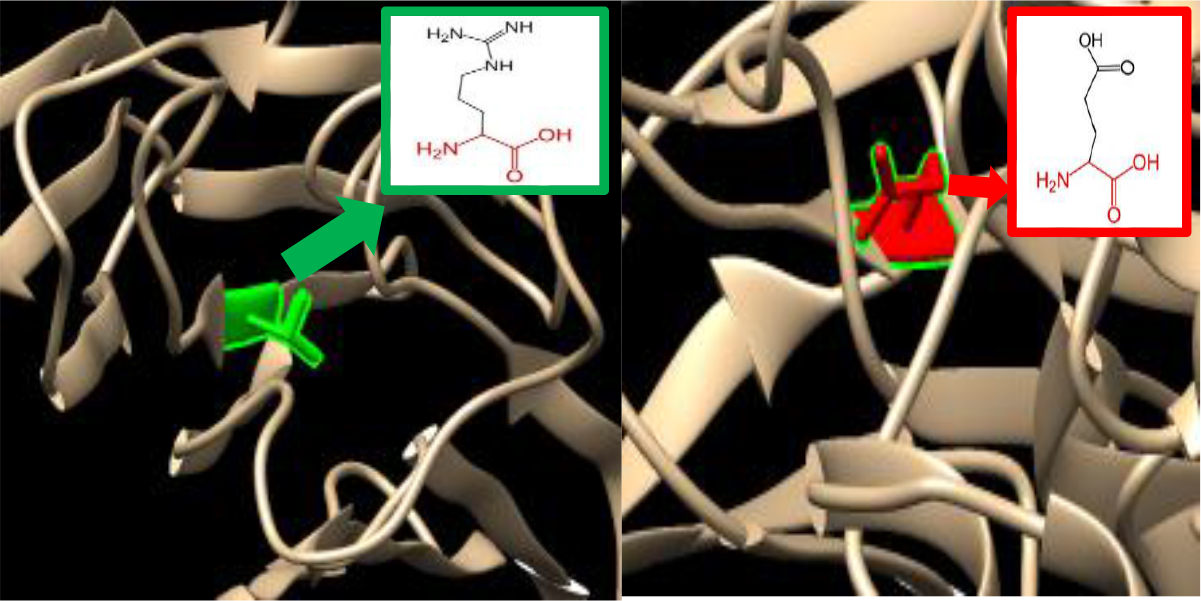
(rs1039834492): (V209E): The amino acid Valine change to Glutamic acid at position 209.

**Figure 39:**
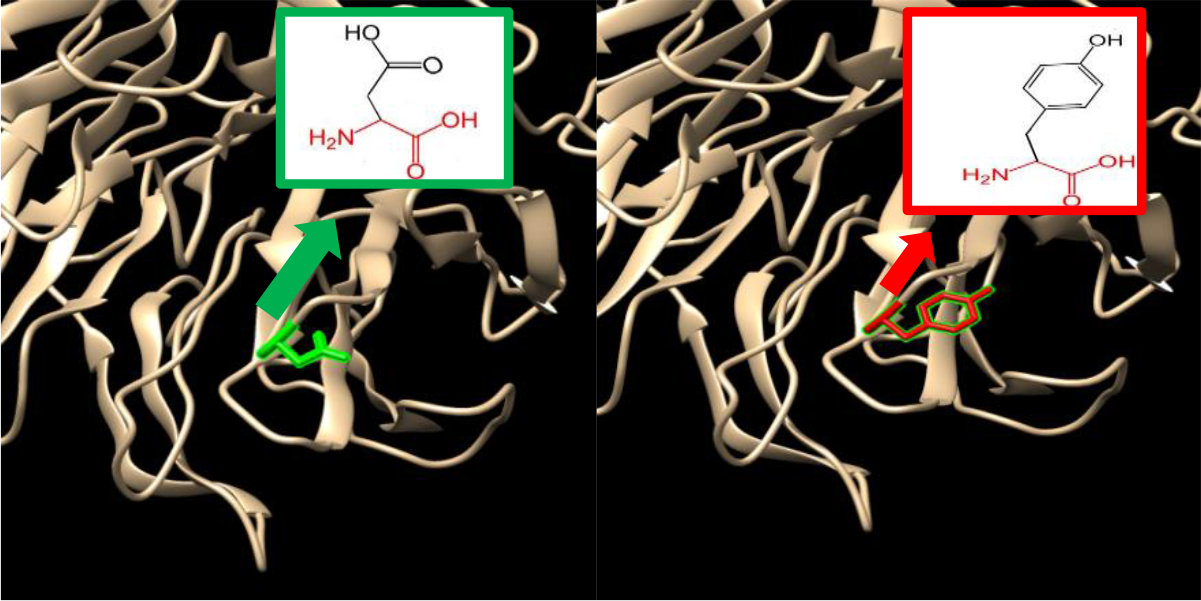
(rs747349523): (D203Y): The amino acid Aspartic acid change to tyrosine at position 203.

**Figure 40:**
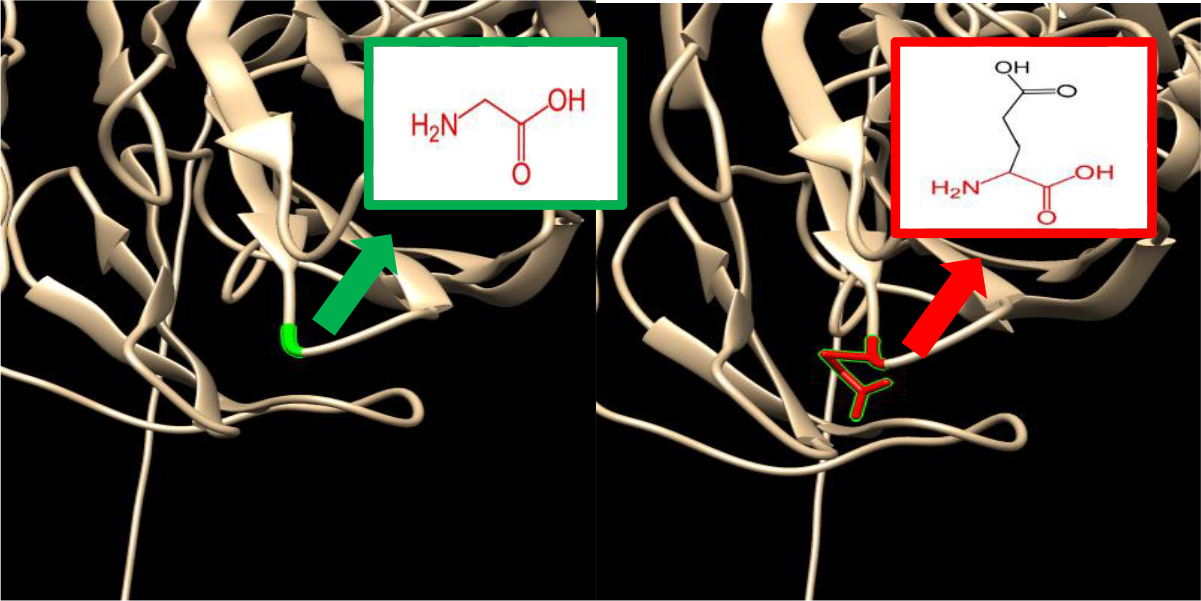
(rs776300430): (G202E): The amino acid Glycine change to Glutamic acid at position 202, Figure was done by chimera 1.8.

**Figure 41:**
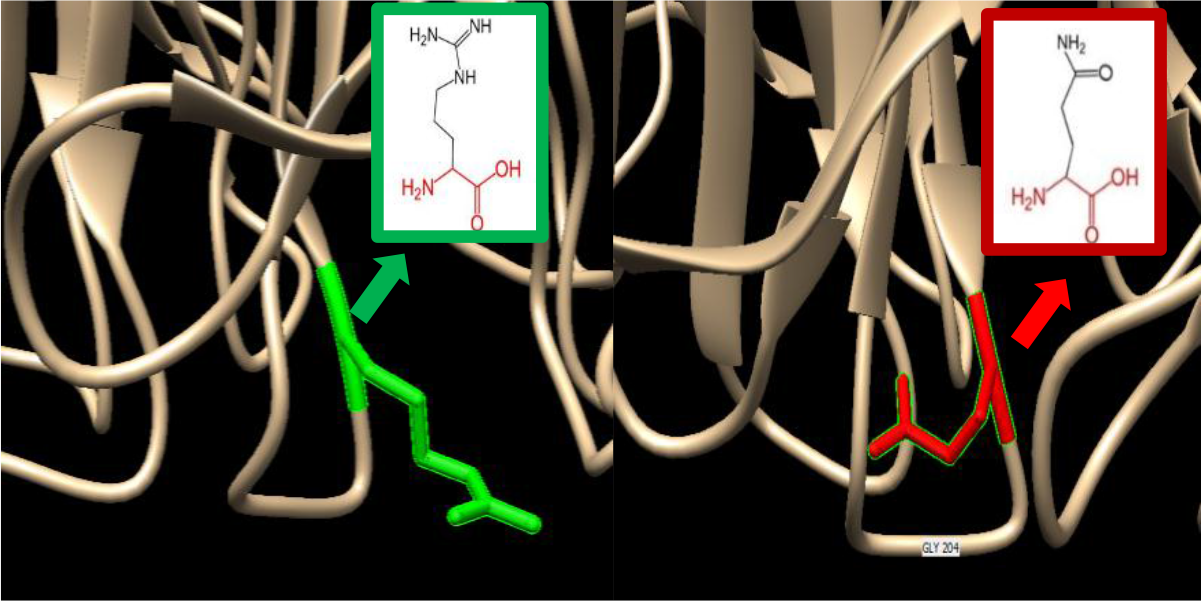
(rs367552387): (R201Q): The amino acid Arginine change to Glutamine at position 201.

**Figure 42:**
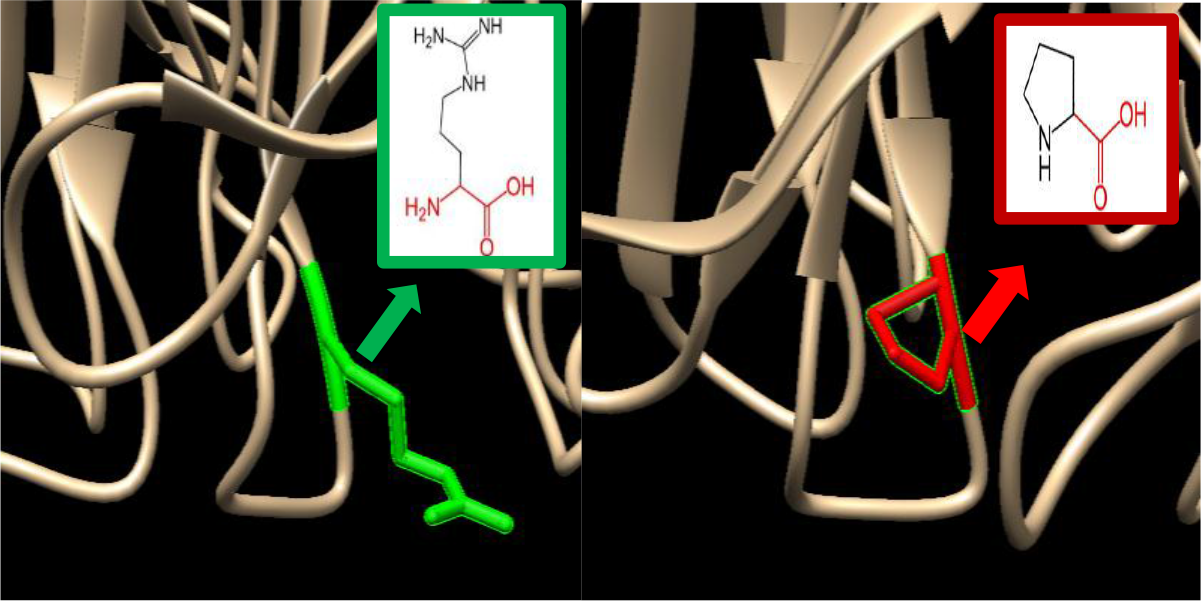
(R201P): The amino acid Arginine change to Proline at position 201.

**Figure 43:**
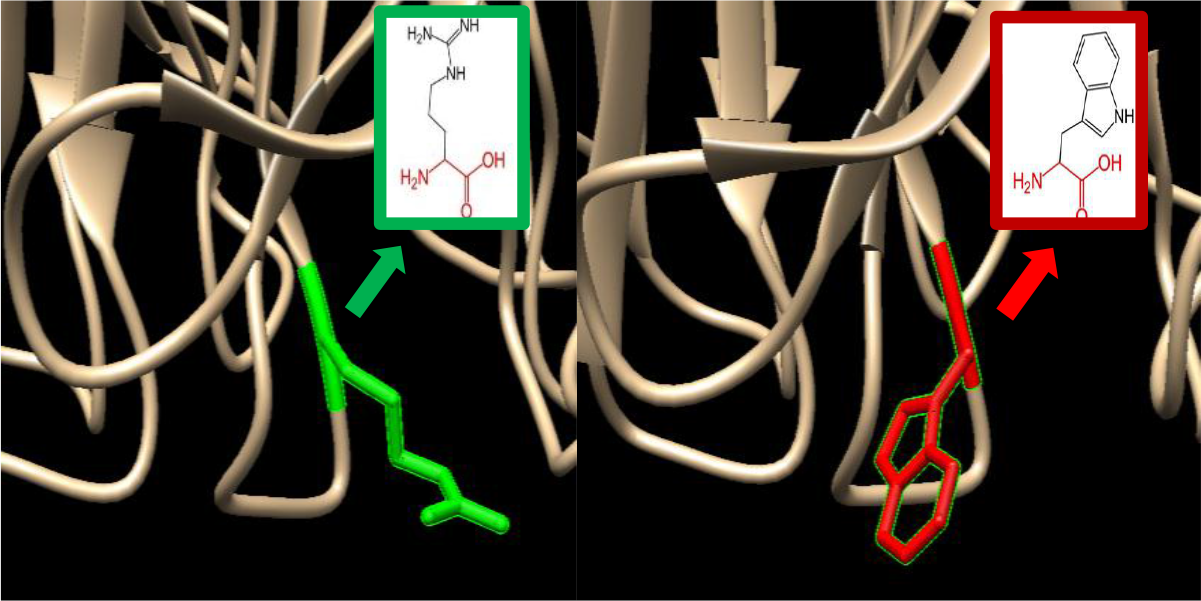
(rs773117166): (R201W): The amino acid Arginine change to Tryptophan at position 201.

**Figure 44:**
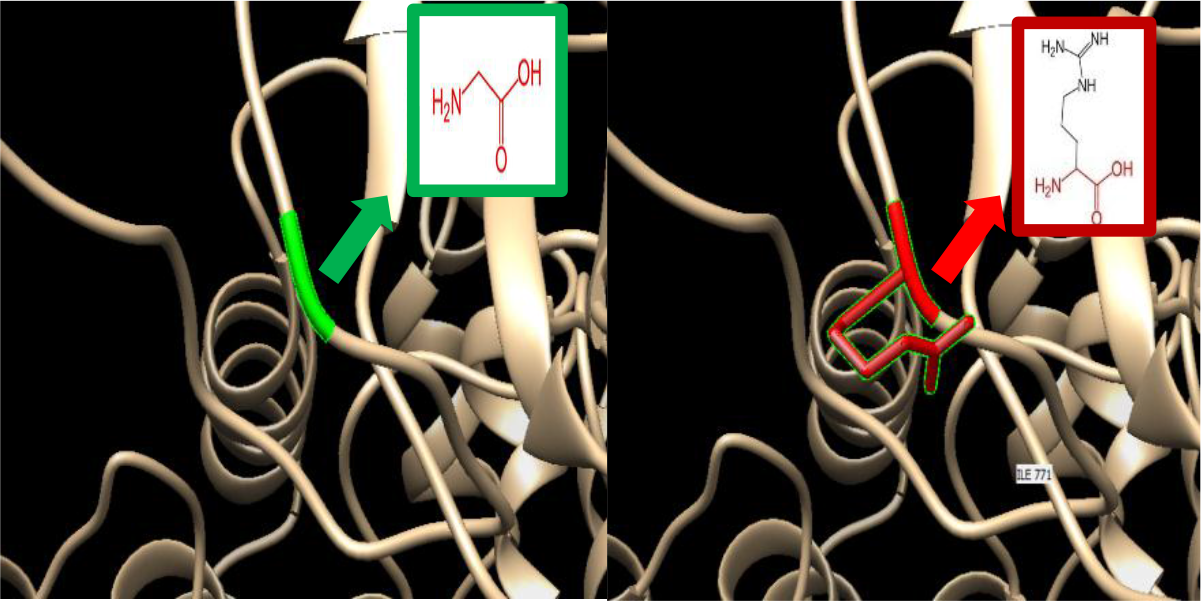
(rs565257577): (G162R): The amino acid Glycine change to Arginine at position 162.

We also used Project HOPE server to submit the 41 most deleterious (damaging) nsSNPs: (rs377679012):(R1137C):Cysteine (The mutant residue) is smaller than Arginine (the wild-type residue.) The wild-type residue charge was POSITIVE; the mutant residue charge is NEUTRAL. The mutant residue is more hydrophobic than the wild-type residue there is a difference in charge between the wild-type and mutant amino acid. The charge of the wild-type residue will be lost; this can cause loss of interactions with other molecules or residues. The wild-type and mutant amino acids differ in size. The mutant residue is smaller; this might lead to loss of interactions. The hydrophobicity of the wild-type and mutant residue differs . The mutation introduces a more hydrophobic residue at this position. This can result in loss of hydrogen bonds and/or disturb correct folding.

(rs372723869):(Y992C): Cysteine (the mutant residue) is smaller than the Tyrosine (wild-type residue). The mutant residue is more hydrophobic than the wild-type residue . The wild-type residue is very conserved, but a few other residue types have been observed at this position too . The mutant residue was not among the other residue types observed at this position in other, homologous proteins. The wild-type and mutant amino acids differ in size. The mutant residue is smaller than the wild-type residue . The mutation will cause an empty space in the core of the protein. The hydrophobicity of the wild-type and mutant residue differs . The mutation will cause loss of hydrogen bonds in the core of the protein and as a result disturb correct folding.

(rs924119518):(Y943C): Cysteine (the mutant residue) is smaller than Tyrosine (the wild-type residue) the mutant residue is more hydrophobic than the wild-type residue. The wild-type residue is very conserved, but a few other residue types have been observed at this position too. The mutant residue was not among the other residue types observed at this position in other, homologous proteins and is located near a highly conserved position. The wild-type and mutant amino acids differ in size. The mutant residue is smaller than the wild-type residue. This will cause a possible loss of external interactions . The hydrophobicity of the wild-type and mutant residue differs.

(rs1319436059):(Y943N): Asparagine (the mutant residue) is smaller than Tyrosine (the wild-type residue). The wild-type residue is more hydrophobic than the mutant residue and it is very conserved, but a few other residue types have been observed at this position too. The mutant residue was not among the other residue types observed at this position in other, homologous proteins and it is located near a highly conserved position. The wild-type and mutant amino acids differ in size. The mutant residue is smaller than the wild-type residue. This will cause a possible loss of external interactions. The hydrophobicity of the wild-type and mutant residue differs. The mutation might cause loss of hydrophobic interactions with other molecules on the surface of the protein.

(rs199617267):(I936T):Threonine(The mutant residue) is smaller than the Isoleucine (wild-type residue). The wild-type residue is more hydrophobic than the mutant residue . The wild-type and mutant amino acids differ in size. The mutant residue is smaller than the wild-type residue. The mutation will cause an empty space in the core of the protein. The mutation will cause loss of hydrophobic interactions in the core of the protein.

(rs1218321803):(A907T): Threonine (The mutant residue) is bigger than Alanine (the wild-type residue). The wild-type residue is more hydrophobic than the mutant residue. The mutation converts the wild-type residue in a residue that does not prefer α-helices as secondary structure. The wild-type residue is very conserved, but a few other residue types have been observed at this position too. The wild-type and mutant amino acids differ in size. The mutant residue is bigger than the wild-type residue. The wild-type residue was buried in the core of the protein. The mutant residue is bigger and probably will not fit. The hydrophobicity of the wild-type and mutant residue differs. The mutation will cause loss of hydrophobic interactions in the core of the protein.

(rs199679232):(Y898C): Cysteine (The mutant residue) is smaller than (the wild-type residue). The mutant residue is more hydrophobic than the wild-type residue. The size difference between wild-type and mutant residue makes that the new residue is not in the correct position to make the same hydrogen bond as the original wild-type residue did. The difference in hydrophobicity will affect hydrogen bond formation. The wild-type residue is very conserved, but a few other residue types have been observed at this position too. The wild-type and mutant amino acids differ in size. The mutant residue is smaller than the wild-type residue. This will cause a possible loss of external interactions. The hydrophobicity of the wild-type and mutant residue differs.

(rs752349406):(N888Y): Tyrosine (The mutant residue) is bigger than (the wild-type residue). The mutant residue is more hydrophobic than the wild-type residue. The wild-type residue is very conserved, but a few other residue types have been observed at this position too. The mutant residue was not among the other residue types observed at this position in other, homologous proteins. However, residues that have some properties in common with your mutated residue were observed. This means that in some rare cases the mutation might occur without damaging the protein. The mutated residue is located on the surface of a domain with unknown function. The wild-type and mutant amino acids differ in size. The mutant residue is bigger than the wild-type residue. The residue is located on the surface of the protein; mutation of this residue can disturb interactions with other molecules or other parts of the protein. The hydrophobicity of the wild-type and mutant residue differs.

(rs1270126477);(D886V): Valine (The mutant residue) is smaller than Aspartic Acid (the wild-type residue). The wild-type residue charge was NEGATIVE; the mutant residue charge is NEUTRAL. The mutant residue is more hydrophobic than the wild-type residue. The size difference between wild-type and mutant residue makes that the new residue is not in the correct position to make the same hydrogen bond as the original wild-type residue did. The difference in hydrophobicity will affect hydrogen bond formation. The mutated residue is located on the surface of a domain with unknown function. There is a difference in charge between the wild-type and mutant amino acid. The charge of the wild-type residue is lost by this mutation. This can cause loss of interactions with other molecules. The wild-type and mutant amino acids differ in size. The mutant residue is smaller than the wild-type residue. This will cause a possible loss of external interactions. The hydrophobicity of the wild-type and mutant residue differs.

(rs1181862976):(Y797C): cysteine (The mutant residue) is smaller than tyrosine (the wild-type residue). Each amino acid has it is own specific size charge, and hydrophobicity-value. The original wild-type residue and newly introduced mutant residue often differ in the properties . The mutant residue is smaller than the wild-type residue. The mutant residue is more hydrophobic than the wild-type residue.

(rs201596987):(T793R): Arginine (The mutant residue) is bigger than Threonine (The wile-type residue). The wild-type residue is very conserved, but a few other residue types have been observed at this position too. Is buried in the core of a domain. The differences between the wild-type and mutant residue might disturb the core structure of this domain. There is a difference in charge between the wild-type and mutant amino acid. The mutant residue introduces a charge in a buried residue which can lead to protein folding problems. The wild-type and mutant amino acids differ in size. The mutant residue is bigger than wild-type residue.

The wild-type residue was buried in the core of the protein. The mutant residue is bigger and probably will not fit. The hydrophobicity of the wild-type and mutant residue differ. The mutation will cause loss of hydrophobic interaction in the core of the protein.

(D791V): Valine (The mutant residue) is smaller than Aspartic Acid (The wild-type residue). The wild-type residue is very conserved, but a few other residue types have been observed at this position too. The mutant residue was not among the other residue types observed at this position in other, homologous proteins. However, the residues that have in some rare cases the mutation might occur without damaging the protein. The mutant residue is located on the surface of a domain with unknown function. The residue was not found to be in contact with other domains of which the function is known without the used structure. There is a difference in charge between the wild-type and mutant amino acid. The wild-type and mutant amino acids differ in size. The mutant reside is smaller than wild-type residue. This will cause a possible loss of external interactions. The hydrophobicity of the wild-type and mutant reside differs.

(rs1452809051):(R749K): Lysine (The mutant residue) is smaller than Arginine (wild-type residue). The wild-type residue is very conserved, but a few other residue types have been observed at this position too. The mutant residue was not among the other residue types observed at this position in other, homologous proteins. Only this type was found at this position. Mutation of a 100% conserved residue is usually damaging for the protein. This mutation might occur in some rare cases, but it is more likely that the mutation is damaging to the protein. The mutated residue is located on the surface of a domain with unknown function. The residue was not found to be in contact with other domains of which the function is known within the used structure. However, contact with other molecules or domains is still possible and might be affected this mutation. The wild-type and mutant amino acid differ in size. The mutant residue is smaller than the wild-type residue. This will cause a possible loss of external interactions.

(rs370575901):(R722Q): Glutamine (The mutant residue) is smaller than Arginine (wild-type residue). The wild-type residue charge was POSITIVE. The mutant residue charge is NATURAL. Only this residue type was found at this position. Mutation of a 100% conserved residue is usually damaging for the protein. This mutation might occur in some rare cases, but it is more likely that the mutation is damaging to the protein. The mutant residue is located near a highly conserved position. The mutated residue is located on the surface of a domain with unknown function. The residue was not found to be in contact with other domains of which the function is known within the used structure. There is a difference charge between the wild-type and mutant amino acid. The charge of the wild-type residue is lost by this mutation. This can cause loss of interactions. The wild-type and mutant amino acids differ in size. The mutant residue is smaller than the wild-type residue. This will cause a possible loss of external interactions.

(rs767003999): (L716S): Serine (The mutant residue) is smaller than Leucine (The wild-type residue). The mutation introduces an amino acid with different properties, which can disrupt this domain and abolish its function. The mutant residue is less hydrophobic than the wild-type residue. Hydrophobic interactions, either in the core of the protein or on the surface, will be lost.

(rs1275065654): (Q710R): Arginine (The mutant residue) is bigger than Glutamine (The wild-type residue). The increases in the size of the mutant residue might lead to bumps. The mutant residue is positively charge, while the wild-type residue charge is neutral. This charge changes could lead to repulsion of ligands or other residues with the same charge.

(rs763692410): (R693W): Tryptophan (The mutant residue) is bigger than Arginine (The wild-type residue). The increases in the size of the mutant residue might lead to bumps. The mutant residue charge is neutral, while the wild-type residue is positively charged. This change can cause loss of interactions with other molecules or residues. The mutant residue is more hydrophobic than the wild-type residue. This can result in loss of hydrogen bonds and/or disturb correct folding.

(rs201742754): (R689Q): Glutamine (The mutant residue) is smaller than Arginine (The wild-type residue). This might lead to loss of interactions. The mutant residue is neutral in charge, while the wild-type residue is positively charged. The loss of this charge can cause loss of interactions with other molecules or residues.

(rs201390288): (R689W): Tryptophan (The mutant residue) is bigger than Arginine (The wild-type residue). The increases in the size of the mutant residue might lead to bumps. The mutant residue charge is neutral, while the wild-type residue is positively charged. This change can cause loss of interactions with other molecules or residues. The mutant residue is more hydrophobic than the wild-type residue. This can result in loss of hydrogen bonds and/or disturb correct folding.

(rs56229130):(G576D): Aspartic Acid (the mutant residue) is bigger than glycine (the wild type). The wild-type residue charge was neutral, the mutant residue charge is negative. The wild-type residue is more hydrophobic than the mutant residue. The wild type residue is the most flexible of all residues. This flexibility might be necessary for the protein’s function. Mutation of this glycine can abolish this function. There is a difference in charge between the wild type and mutant amino acid. The mutation introduces a charge; this can cause repulsion of ligands or other residues with the same charge. The wild type and mutant amino acids differ in size. The mutant residue is bigger, this might lead to bumps. The torsion angles for this residue are unusual. Only glycine is flexible enough to make these torsion angles, mutation into another residue will force the local backbone into an incorrect conformation and will disturb the local structure.

(rs375957332):(P410L): Leucine (The mutant residue) is bigger than the Proline (The wild type). Proline are known to be very rigid and therefore induce a special backbone conformation, which might be required at this position. The mutation can disturb this special conformation, there by disturbing the local structure. The wild type and mutant amino acids differ in size. The mutant residue is bigger this might lead to bumps.

(rs1414061926):(G396V): Valine (the mutant residue) is bigger than Glycine (the wild type)the mutant residue is more hydrophobic than the wild-type residue. The torsion angles for this residue are unusual. Only glycine is flexible enough to make these torsion angles, mutation into another residue will force the local backbone into an incorrect conformation and will disturb the local structure. The wild type and mutant amino acids differ in size. The mutant residue is bigger this might lead to bumps.

(rs775321513):(L336S): Serine (the mutant residue) is smaller than leucine (the wild type residue). The wild type residue is more hydrophobic than the mutant residue. The hydrophobicity of the wild type and mutant residue differs, so hydrophobic interactions, either in the core of the protein or on the surface, will be lost. The wild type and mutant amino acids differ in size. The mutant residue is smaller; this might lead to loss of interactions.

(rs764033545):(H329Q): Histidine (the mutant residue) is smaller than glutamine (the wild type),the wild-type and mutant amino acids differ in size. The mutant residue is smaller; this might lead to loss of interactions.

(rs750663365):(Y328S): Serine (The mutant residue) is smaller than Tyrosine (the wild-type residue). The wild-type residue is predicted (using the Reprof software) to be located in its preferred secondary structure, a β-strand. The mutant residue prefers to be in another secondary structure; therefore the local conformation will be slightly destabilized. The wild-type and mutant amino acids differ in size. The mutant residue is smaller; this might lead to loss of interactions.

(Y328C): cysteine (The mutant residue) is smaller than (the wild-type residue). The mutant residue is more hydrophobic than the wild-type residue. The wild-type residue is predicted (using the Reprof software) to be located in its preferred secondary structure, a β-strand. The mutant residue prefers to be in another secondary structure; therefore the local conformation will be slightly destabilized. The wild-type and mutant amino acids differ in size. The mutant residue is smaller; this might lead to loss of interactions. The hydrophobicity of the wild-type and mutant residue differs. The mutation introduces a more hydrophobic residue at this position. This can result in loss of hydrogen bonds and/or disturb correct folding.

(Y328F): Phenylalanine (The mutant residue) is bigger than Tyrosine (the wild-type residue). The mutant residue is more hydrophobic than the wild-type residue. The wild-type and mutant amino acids differ in size. The mutant residue is smaller; this might lead to loss of interactions. The hydrophobicity of the wild-type and mutant residue differs. The mutation introduces a more hydrophobic residue at this position. This can result in loss of hydrogen bonds and/or disturb correct folding.

(rs1159235595):(W323R): Arginine (The mutant residue) is smaller than Tryptophan (the wild-type residue). The wild-type residue charge was NEUTRAL, the mutant residue charge is POSITIVE. The wild-type residue is predicted (using the Reprof software) to be located in its preferred secondary structure, a β-strand. The mutant residue prefers to be in another secondary structure; therefore the local conformation will be slightly destabilized. The wild-type residue is more hydrophobic than the mutant residue. The wild-type and mutant amino acids differ in size. The mutant residue is smaller; this might lead to loss of interactions. The hydrophobicity of the wild-type and mutant residue differs. Hydrophobic interactions, either in the core of the protein or on the surface, will be lost.

(rs1288706972):(S298F): Phenylalanine (The mutant residue) is bigger than Serine (the wild-type residue). The mutant residue is more hydrophobic than the wild-type residue. The wild-type residue is predicted (using the Reprof software) to be located in its preferred secondary structure, a turn. The mutant residue prefers to be in another secondary structure; therefore the local conformation will be slightly destabilized. The wild-type and mutant amino acids differ in size. The mutant residue is bigger, this might lead to bumps. The hydrophobicity of the wild-type and mutant residue differs. The mutation introduces a more hydrophobic residue at this position. This can result in loss of hydrogen bonds and/or disturb correct folding.

()(G272R) Arginine (The mutant residue) is bigger than Glycine (the wild-type residue). The wild-type residue charge was NEUTRAL, the mutant residue charge is POSITIVE. The wild-type residue is more hydrophobic than the mutant residue. There is a difference in charge between the wild-type and mutant amino acid. The mutation introduces a charge, this can cause repulsion of ligands or other residues with the same charge. The wild-type and mutant amino acids differ in size. The mutant residue is bigger, this might lead to bumps. The torsion angles for this residue are unusual. only glycine is flexible enough to make these torsion angles, mutation into another residue will force the local backbone into an incorrect conformation and will disturb the local structure.

(rs1394468770):(P248L): Leucine (The mutant residue) is bigger than Proline (The wild-type residue). The mutated residue is located in a domain that is important for binding of other molecules. Mutation of the residue might disturb this function. The wild-type and mutant amino acids differ in size. The mutant residue is bigger, this might lead to bumps. Prolines are known to have a very rigid structure, sometimes forcing the backbone in a specific conformation. Possibly, the mutation changes a proline with such a function into another residue, thereby disturbing the local structure.

(rs1312638005):(V223E): Glutamic acid (The mutant residue) is bigger than Valine (The wild-type residue). The wild-type residue charge was NEUTRAL; the mutant residue charge is NEGATIVE. The wild-type residue is more hydrophobic than the mutant residue. There is a difference in charge between the wild-type and mutant amino acid. The mutation introduces a charge, this can cause repulsion of ligands or other residues with the same charge. The wild-type and mutant amino acids differ in size. The mutant residue is bigger, this might lead to bumps. The hydrophobicity of the wild-type and mutant residue differs. Hydrophobic interactions, either in the core of the protein or on the surface, will be lost.

(rs749382362):(R219Q): Glutamine (The mutant residue) is smaller than Arginine (The wild-type residue). The wild-type residue charge was POSITIVE; the mutant residue charge is NEUTRAL. The mutated residue is located in a domain that is important for binding of other molecules. Mutation of the residue might disturb this function. There is a difference in charge between the wild-type and mutant amino acid. The charge of the wild-type residue will be lost; this can cause loss of interactions with other molecules or residues. The wild-type and mutant amino acids differ in size. The mutant residue is smaller; this might lead to loss of interactions.

(rs374238430):(R219W): Tryptophan (The mutant residue) is bigger than Arginine (The wild-type residue). The wild-type residue charge was POSITIVE; the mutant residue charge is NEUTRAL. The mutant residue is more hydrophobic than the wild-type residue. The mutated residue is located in a domain that is important for binding of other molecules. Mutation of the residue might disturb this function. There is a difference in charge between the wild-type and mutant amino acid. The charge of the wild-type residue will be lost; this can cause loss of interactions with other molecules or residues. The wild-type and mutant amino acids differ in size. The mutant residue is bigger, this might lead to bumps. The hydrophobicity of the wild-type and mutant residue differs. The mutation introduces a more hydrophobic residue at this position. This can result in loss of hydrogen bonds and/or disturb correct folding.

(rs1039834492):(V209E): Glutamic Acid (The mutant residue) is bigger than Valine (The wild-type residue). The wild-type residue charge was NEUTRAL; the mutant residue charge is NEGATIVE. The wild-type residue is more hydrophobic than the mutant residue. The mutated residue is located in a domain that is important for binding of other molecules. Mutation of the residue might disturb this function. There is a difference in charge between the wild-type and mutant amino acid. The mutation introduces a charge; this can cause repulsion of ligands or other residues with the same charge. The wild-type and mutant amino acids differ in size. The mutant residue is bigger, this might lead to bumps. The hydrophobicity of the wild-type and mutant residue differs. Hydrophobic interactions, either in the core of the protein or on the surface, will be lost.

(rs747349523):(D203Y): Tyrosine (The mutant residue) is bigger than Aspartic Acid (the wild-type residue). The wild-type residue charge was NEGATIVE, the mutant residue charge is NEUTRAL . The mutant residue is more hydrophobic than the wild-type residue There is a difference in charge between the wild-type and mutant amino acid. The charge of the wild-type residue will be lost, this can cause loss of interactions with other molecules or residues. The wild-type and mutant amino acids differ in size . The mutant residue is bigger, this might lead to bumps . The hydrophobicity of the wild-type and mutant residue differs . The mutation introduces a more hydrophobic residue at this position. This can result in loss of hydrogen bonds and/or disturb correct folding.

(rs776300430):(G202E): Glutamic Acid (The mutant residue) is bigger than Glycine( the wild-type residue). The wild-type residue charge was NEUTRAL, the mutant residue charge is NEGATIVE. The wild-type residue is more hydrophobic than the mutant residue there is a difference in charge between the wild-type and mutant amino acid. The mutation introduces a charge; this can cause repulsion of ligands or other residues with the same charge. The wild-type and mutant amino acids differ in size. The mutant residue is bigger, this might lead to bump. The torsion angles for this residue are unusual. Only glycine is flexible enough to make these torsion angles, mutation into another residue will force the local backbone into an incorrect conformation and will disturb the local structure.

(rs367552387):(R201Q): Glutamine (The mutant residue) is smaller than Arginine (the wild-type residue). The wild-type residue charge was POSITIVE, the mutant residue charge is NEUTRAL. There is a difference in charge between the wild-type and mutant amino acid. The charge of the wild-type residue will be lost, this can cause loss of interactions with other molecules or residues. The wild-type and mutant amino acids differ in size . The mutant residue is smaller, this might lead to loss of interactions.

()(R201P): Proline (The mutant residue) is smaller than Arginine (the wild-type residue). The wild-type residue charge was POSITIVE, the mutant residue charge is NEUTRAL. The mutant residue is more hydrophobic than the wild-type residue There is a difference in charge between the wild-type and mutant amino acid. The charge of the wild-type residue will be lost, this can cause loss of interactions with other molecules or residues. The wild-type and mutant amino acids differ in size. The mutant residue is smaller, this might lead to loss of interactions. The hydrophobicity of the wild-type and mutant residue differs. The mutation introduces a more hydrophobic residue at this position. This can result in loss of hydrogen bonds and/or disturb correct folding.

(rs773117166)(R201W) Tryptophan (The mutant residue) is bigger than Arginine (the wild-type residue.)The wild-type residue charge was POSITIVE, the mutant residue charge is NEUTRAL The mutant residue is more hydrophobic than the wild-type residue There is a difference in charge between the wild-type and mutant amino acid. The charge of the wild-type residue will be lost, this can cause loss of interactions with other molecules or residues. The wild-type and mutant amino acids differ in size. The mutant residue is bigger, this might lead to bumps . The hydrophobicity of the wild-type and mutant residue differs . The mutation introduces a more hydrophobic residue at this position. This can result in loss of hydrogen bonds and/or disturb correct folding.

(rs565257577)(G162R) Arginine (The mutant residue) is bigger than Glycine( the wild-type residue). The wild-type residue charge was NEUTRAL, the mutant residue charge is POSITIVE. The wild-type residue is more hydrophobic than the mutant residue. There is a difference in charge between the wild-type and mutant amino acid. The mutation introduces a charge, this can cause repulsion of ligands or other residues with the same charge. The wild-type and mutant amino acids differ in size. The mutant residue is bigger, this might lead to bumps. The torsion angles for this residue are unusual. only glycine is flexible enough to make these torsion angles, mutation into another residue will force the local backbone into an incorrect conformation and will disturb the local structure.

In the light of this study, any one with relation to Jewish ancestors or suspected of being a half Jewish, should be consider to do Exome sequencing for IKBKAP gene to check Familial dysautonomia disease.

## Conclusion

In the current work the influence of functional SNPs in the *IKBKAP* gene was investigated through various computational methods, and determined that (R1137C, Y992C, Y943C, Y943N, I936T, A907T, Y898C, N888Y, D886V, Y797C, T793R, D791V, R749K, R722Q, L716S, Q710R, R693W, R689Q, R689W, G576D, P410L, G396V, L336S, H329Q, Y328S, Y328C, Y328F, W323R, S298F, G272R, P248L, V223E, R219Q, R219W, V209E, D203Y, G202E, R201Q, R201P, R201W, G162R) new SNPs have a potential functional impact and can thus be used as diagnostic markers.

## Acknowledgment

The authors wish to express their profound gratitude and deep regards to Africa City of Technology - Sudan.

## Data availability statement

Data available on request from the authors, the data that support the findings of this study are available from the cross ponding author upon reasonable request.

## References

1. Axelrod FB. Familial dysautonomia. Muscle & nerve. 2004;29(3):352–63.

2. Axelrod FB, Hilz MJ. Inherited autonomic neuropathies. Seminars in neurology. 2003;23(4):381–90.

3. Donadon I, Pinotti M, Rajkowska K, Pianigiani G, Barbon E, Morini E, et al. Exon-specific U1 snRNAs improve ELP1 exon 20 definition and rescue ELP1 protein expression in a familial dysautonomia mouse model. Human molecular genetics. 2018;27(14):2466–76.

4. Palma JA, Norcliffe-Kaufmann L, Fuente-Mora C, Percival L, Mendoza-Santiesteban C, Kaufmann H. Current treatments in familial dysautonomia. Expert opinion on pharmacotherapy. 2014;15(18):2653–71.

5. Slaugenhaupt SA, Blumenfeld A, Gill SP, Leyne M, Mull J, Cuajungco MP, et al. Tissue-specific expression of a splicing mutation in the IKBKAP gene causes familial dysautonomia. American journal of human genetics. 2001;68(3):598–605.

6. Carroll MS, Kenny AS, Patwari PP, Ramirez JM, Weese-Mayer DE. Respiratory and cardiovascular indicators of autonomic nervous system dysregulation in familial dysautonomia. Pediatric pulmonology. 2012;47(7):682–91.

7. Palma JA, Norcliffe-Kaufmann L, Perez MA, Spalink CL, Kaufmann H. Sudden Unexpected Death During Sleep in Familial Dysautonomia: A Case-Control Study. Sleep. 2017;40(8).

8. Dietrich P, Dragatsis I. Familial Dysautonomia: Mechanisms and Models. Genetics and molecular biology. 2016;39(4):497–514.

9. Dietrich P, Alli S, Shanmugasundaram R, Dragatsis I. IKAP expression levels modulate disease severity in a mouse model of familial dysautonomia. Human molecular genetics. 2012;21(23):5078–90.

10. Di Santo R, Bandau S, Stark MJ. A conserved and essential basic region mediates tRNA binding to the Elp1 subunit of the Saccharomyces cerevisiae Elongator complex. Molecular microbiology. 2014;92(6):1227–42.

11. Xu H, Lin Z, Li F, Diao W, Dong C, Zhou H, et al. Dimerization of elongator protein 1 is essential for Elongator complex assembly. Proceedings of the National Academy of Sciences of the United States of America. 2015;112(34):10697–702.

12. Lehavi O, Aizenstein O, Bercovich D, Pavzner D, Shomrat R, Orr-Urtreger A, et al. Screening for familial dysautonomia in Israel: evidence for higher carrier rate among Polish Ashkenazi Jews. Genetic testing. 2003;7(2):139–42.

13. Sinha R, Kim YJ, Nomakuchi T, Sahashi K, Hua Y, Rigo F, et al. Antisense oligonucleotides correct the familial dysautonomia splicing defect in IKBKAP transgenic mice. Nucleic acids research. 2018;46(10):4833–44.

14. Salani M, Urbina F, Brenner A, Morini E, Shetty R, Gallagher CS, et al. Development of a Screening Platform to Identify Small Molecules That Modify ELP1 Pre-mRNA Splicing in Familial Dysautonomia. SLAS discovery : advancing life sciences R & D. 2018:2472555218792264.

15. Leyne M, Mull J, Gill SP, Cuajungco MP, Oddoux C, Blumenfeld A, et al. Identification of the first non-Jewish mutation in familial Dysautonomia. American journal of medical genetics Part A. 2003;118A(4):305–8.

16. Cuajungco MP, Leyne M, Mull J, Gill SP, Lu W, Zagzag D, et al. Tissue-specific reduction in splicing efficiency of IKBKAP due to the major mutation associated with familial dysautonomia. American journal of human genetics. 2003;72(3):749–58.

17. Kazachkov M, Palma JA, Norcliffe-Kaufmann L, Bar-Aluma BE, Spalink CL, Barnes EP, et al. Respiratory care in familial dysautonomia: Systematic review and expert consensus recommendations. Respiratory medicine. 2018;141:37–46.

18. Norcliffe-Kaufmann L, Axelrod FB, Kaufmann H. Developmental abnormalities, blood pressure variability and renal disease in Riley Day syndrome. Norcliffe-Journal of human hypertension. 2013;27(1):51–5.

19. Mendoza-Santiesteban CE, Hedges ITRii, Norcliffe-Kaufmann L, Axelrod F, Kaufmann H. Selective retinal ganglion cell loss in familial dysautonomia. Journal of neurology. 2014;261(4):702–9.

20. Portnoy S, Maayan C, Tsenter J, Ofran Y, Goldman V, Hiller N, et al. Characteristics of ataxic gait in familial dysautonomia patients. PloS one. 2018;13(4):e0196599.

21. Bajard A, Chabaud S, Cornu C, Castellan AC, Malik S, Kurbatova P, et al. An in silico approach helped to identify the best experimental design, population, and outcome for future randomized clinical trials. Journal of clinical epidemiology. 2016;69:125–36.

22. Benson DA, Cavanaugh M, Clark K, Karsch-Mizrachi I, Lipman DJ, Ostell J, et al. GenBank. Nucleic acids research. 2017;45(D1):D37–d42.

23. Artimo P, Jonnalagedda M, Arnold K, Baratin D, Csardi G, de Castro E, et al. ExPASy: SIB bioinformatics resource portal. Nucleic acids research. 2012;40(Web Server issue):W597–603.

24. Sim NL, Kumar P, Hu J, Henikoff S, Schneider G, Ng PC. SIFT web server: predicting effects of amino acid substitutions on proteins. Nucleic acids research. 2012;40(Web Server issue):W452–7.

25. Adzhubei IA, Schmidt S, Peshkin L, Ramensky VE, Gerasimova A, Bork P, et al. A method and server for predicting damaging missense mutations. Nature methods. 2010;7(4):248–9.

26. Choi Y, Sims GE, Murphy S, Miller JR, Chan AP. Predicting the functional effect of amino acid substitutions and indels. PloS one. 2012;7(10):e46688.

27. Hecht M, Bromberg Y, Rost B. Better prediction of functional effects for sequence variants. BMC genomics. 2015;16 Suppl 8:S1.

28. Calabrese R, Capriotti E, Fariselli P, Martelli PL, Casadio R. Functional annotations improve the predictive score of human disease-related mutations in proteins. Human mutation. 2009;30(8):1237–44.

29. Lopez-Ferrando V, Gazzo A, de la Cruz X, Orozco M, Gelpi JL. PMut: a web-based tool for the annotation of pathological variants on proteins, 2017 update. Nucleic acids research. 2017;45(W1):W222–w8.

30. Capriotti E, Fariselli P, Casadio R. I-Mutant2.0: predicting stability changes upon mutation from the protein sequence or structure. Nucleic acids research. 2005;33(Web Server issue):W306–10.

31. Warde-Farley D, Donaldson SL, Comes O, Zuberi K, Badrawi R, Chao P, et al. The GeneMANIA prediction server: biological network integration for gene prioritization and predicting gene function. Nucleic acids research. 2010;38(Web Server issue):W214–20.

32. Wang S, Li W, Liu S, Xu J. RaptorX-Property: a web server for protein structure property prediction. Nucleic acids research. 2016;44(W1):W430–5.

